# FACT subunit SUPT16H associates with BRD4 and contributes to silencing of antiviral interferon signaling

**DOI:** 10.1101/2021.04.21.440833

**Authors:** Dawei Zhou, Jun-Gyu Park, Zhenyu Wu, Huachao Huang, Guillaume N. Fiches, Ayan Biswas, Tai-Wei Li, Qin Ma, Luis Martinez-Sobrido, Netty Santoso, Jian Zhu

**Affiliations:** Department of Pathology, The Ohio State University Wexner Medical Center, Columbus, OH 43210, USA; Texas Biomedical Research Institute, San Antonio, TX 78227, USA; Department of Medicine, Columbia University Medical Center, New York, NY 10032, USA; Department of Biomedical Informatics, The Ohio State University Wexner Medical Center, Columbus, OH 43210, USA

**Keywords:** FACT complex, SUPT16H, BRD4, TIP60, acetylation, gene silencing, interferon, antiviral immunity, NK cell

## Abstract

FACT (<u>FA</u>cilitates <u>C</u>hromatin <u>T</u>ranscription) is a heterodimeric protein complex composed of SUPT16H and SSRP1, and a histone chaperone participating in chromatin remodeling during gene transcription. FACT complex is profoundly regulated, and contributes to both gene activation and suppression. Here we reported that SUPT16H, a subunit of FACT, is acetylated at lysine 674 (K674) of middle domain (MD), which involves TIP60 histone acetyltransferase. Such acetylation of SUPT16H is recognized by bromodomain protein BRD4, which promotes protein stability of SUPT16H. We further demonstrated that SUPT16H-BRD4 associates with histone modification enzymes (EZH2, HDAC1) and affects histone marks (H3K9me3, H3K27me3 and H3ac). BRD4 is known to profoundly regulate interferon (IFN) signaling, while such function of SUPT16H has never been explored. Surprisingly, our results revealed that SUPT16H genetic knockdown via RNAi or pharmacological inhibition by using its inhibitor, curaxin 137 (CBL0137), results in the induction of IFNs and interferon-stimulated genes (ISGs). Through this mechanism, CBL0137 is shown to efficiently inhibit infection of multiple viruses, including Zika, influenza, and SARS-CoV-2. Furthermore, we demonstrated that CBL0137 also causes the remarkable activation of IFN signaling in natural killer (NK) cells, which promotes the NK-mediated killing of virus-infected cells in a co-culture system using human primary NK cells. Overall, our studies unraveled the previously un-appreciated role of FACT complex in regulating IFN signaling in both epithelial and NK cells, and also proposed the novel application of CBL0137 to treat viral infections.

## Introduction

In eukaryotes, histone chaperones play a critical role in regulating gene expression by maintaining nucleosome assembly and genome stability (Hammond et al., 2017). As one of key histone chaperones, the main function of FACT (<u>FA</u>cilitates <u>C</u>hromatin <u>T</u>ranscription) complex is the deposition and reorganization of histone H2A-H2B and H3-H4 octamers (Belotserkovskaya et al., 2003; Orphanides et al., 1998; Stuwe et al., 2008). FACT participates in various processes, such as gene transcription, DNA replication and repair, and centromere activation (Formosa and Winston, 2020; Murawska and Ladurner, 2016; Orphanides et al., 1998). FACT is heterodimer composed of two subunits, SUPT16H and SSRP1, which interact with each other as well as the nucleosome through specific domains. Beyond the common functions conveyed through FACT, both SUPT16H and SSRP1 possess their additional ones independent from each other (Ding et al., 2016; Wienholz et al., 2019).

Since the initial identification, FACT has been generally believed to facilitate gene transcription due to its control of nucleosome remodeling critical for the RNA polymerase II (Pol II) elongation (Belotserkovskaya et al., 2003). However, there is also increasing evidence supporting that FACT contributes to gene suppression through various mechanisms. For example, earlier studies from us revealed that FACT represses gene expression of HIV-1 proviruses integrated into host genomes and promotes viral latency by interfering with the interaction between P-TEFb and viral Tat-LTR axis (Huang et al., 2015). Likewise, FACT has been recently reported to block lytic reactivation of latent Epstein-Barr virus (EBV) that tethers viral episomes with host genomes by supporting MYC expression in Burkitt lymphoma (BL) (Guo et al., 2020). FACT is known to inhibit the expression of cryptic genes, antisense transcripts, and subtelomeric genes (Kaplan et al., 2003; Murawska et al., 2020). Retro-transposable elements and the associated cryptic promoters are also silenced by FACT (Chen et al., 2020). These data suggest that FACT plays a more profound role in regulating gene expression, while the underlying mechanisms contributing to such functions of FACT need more careful investigations.

One direction is to determine the impact of post-translational modifications (PTMs) on FACT’s transcriptional activities. There has been a study showing that FACT undergoes K63-linked ubiquitination that affects its function during DNA replication (Han et al., 2010). Beyond ubiquitination, other PTMs of FACT proteins are far from detailed characterizations. For example. impact of acetylation on FACT has never been deliberately examined, although several proteomic screenings have predicted that there are multiple acetylation sites of FACT subunit SUPT16H (Beli et al., 2012; Choudhary et al., 2009; Mertins et al., 2013; Weinert et al., 2013). However, there is currently no knowledge regarding whether acetylation of FACT proteins truly exists nor what is the functional contribution of FACT acetylation. Our studies started from the point to characterize the acetylation of FACT proteins, which led to the further findings that FACT subunit SUPT16H associates with BRD4 through its acetylation and contributes to silencing of antiviral interferon (IFN) signaling.

## Results

### SUPT16H is acetylated by TIP60 and interacts with BRD4

To determine the acetylation of FACT proteins, we performed the protein immunoprecipitation (IP) using an anti-acetyl lysine antibody in different cell lines, including HEK293T, HeLa, Jurkat and NK-92 cells, followed by the protein immunoblotting of FACT proteins. Acetylation of SUPT16H, but not SSRP1, can be readily detected in all tested cell lines (**Fig 1A**). We further spent effort searching for the acetyltransferase(s) that contributes to SUPT16H acetylation. We showed that TIP60, a well-studied histone acetyltransferase (HAT) catalyzing acetylation of various histone and nonhistone proteins, interacts with SUPT16H (**Fig 1B**) and that its knockdown by shRNA significantly reduces SUPT16H acetylation (**Fig 1C, D**). Consistently, treatment of a TIP60-specific inhibitor MG149 also led to the reduction of SUPT16H acetylation without obvious cytotoxicity (**Fig 1E, S1A**). Protein acetylation can be recognized by certain readers that further recruit downstream executers to fulfill regulatory functions (Gong et al., 2016). We confirmed that SUPT16H interacts with BRD4, a key acetylation reader containing two bromodomains (**Fig 1F**). Treatment of a bromodomain and extra-terminal motif inhibitor (BETi) JQ1 led to the drastic reduction of SUPT16H and BRD4 interaction (**Fig 1G**), reassuring that BRD4 interaction with SUTP16H is through its acetyl lysine-binding capacity. Furthermore, knockdown of TIP60 also greatly decreased the SUPT16H and BRD4 interaction (**Fig 1H**). Overall, these results indicated that SUPT16H is acetylated by TIP60 and that SUPT16H acetylation mediates protein interaction of SUPT16H and BRD4.

**Fig 1.**
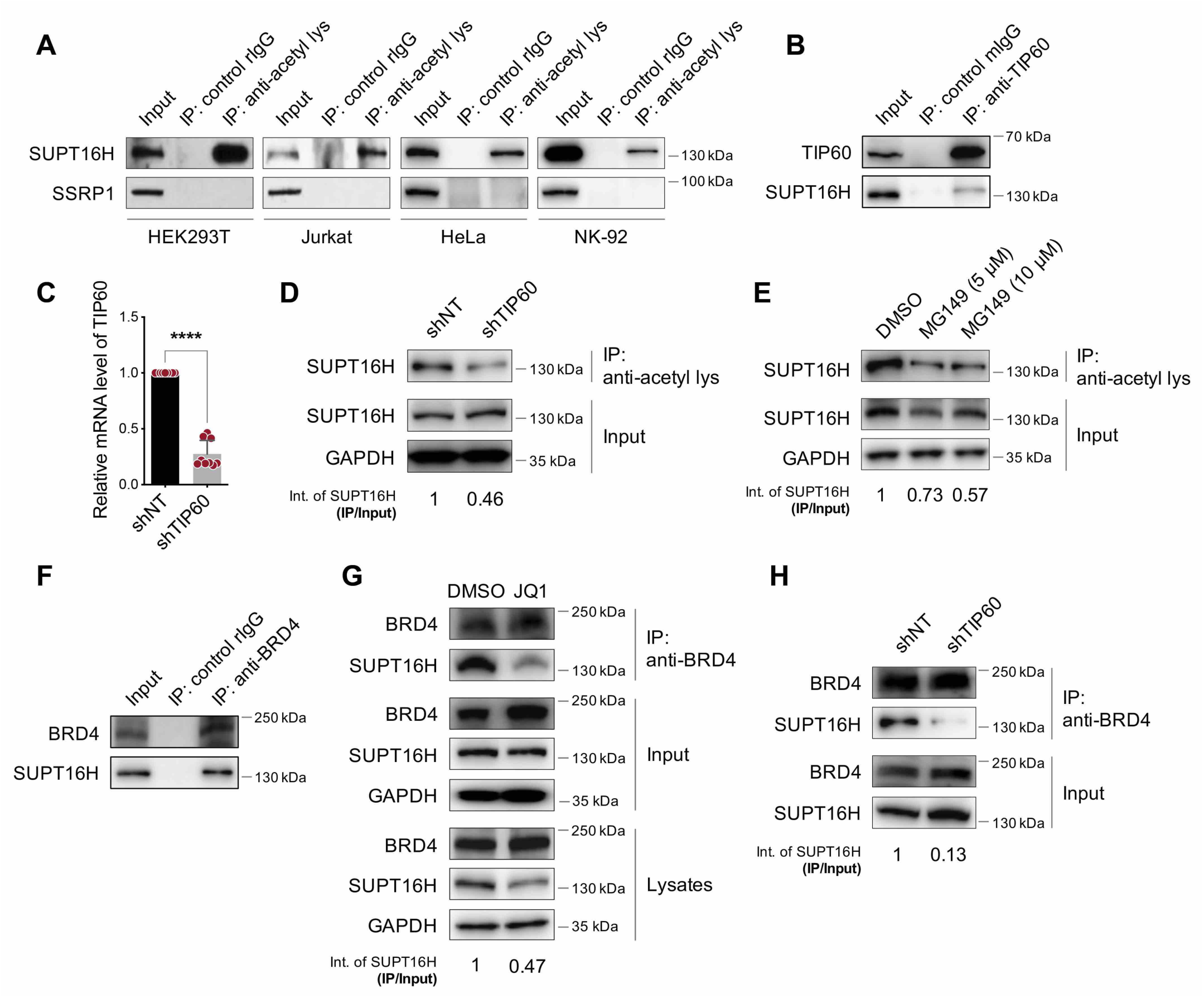
SUPT16H acetylation requires TIP60 and is recognized by BRD4. (**A**) Lysates of HEK293T, Jurkat, HeLa, and NK-92 cells were incubated with an acetyl-lysine antibody or normal mouse IgG (mIgG). Immunoprecipitated protein samples were analyzed by protein immunoblotting using a SUPT16H or SSRP1 antibody. (**B**) Lysates of HEK293T cells were incubated with a TIP60 antibody or mIgG. Immunoprecipitated protein samples were analyzed by protein immunoblotting using a SUPT16H or TIP60 antibody. (**C**) HEK293T cells stably expressing TIP60 or non-targeting (NT) shRNA were subjected to RT-qPCR analysis of TIP60 expression. Results were calculated from three independent experiments (**** *P* < 0.0001, Student’s *t*-test). (**D**) Lysates of HEK293T cells in (**C**) were incubated with an acetyl-lysine antibody. Immunoprecipitated protein samples were analyzed by protein immunoblotting using a SUPT16H antibody. (**E**) Lysates of HEK293T cells treated with a TIP60-specific inhibitor MG149 were incubated with an acetyl-lysine antibody. Immunoprecipitated protein samples were analyzed by protein immunoblotting using a SUPT16H antibody. (**F**) Lysates of HEK293T cells were incubated with a BRD4 antibody or normal rabbit IgG (rIgG). Immunoprecipitated protein samples were analyzed by protein immunoblotting using a SUPT16H or BRD4 antibody. (**G**) Lysates of HEK293T cells treated with a BETi JQ1 were adjusted to assure the same input of SUPT16H, and incubated with a BRD4 antibody. Immunoprecipitated protein samples were analyzed by protein immunoblotting using a SUPT16H or BRD4 antibody. (**H**) Lysates of HEK293T cells in (**C**) were incubated with a BRD4 antibody. Immunoprecipitated protein samples were analyzed by protein immunoblotting using a SUPT16H or BRD4 antibody. (Int: intensity)

### SUPT16H is acetylated at K674 of the middle domain (MD)

We further characterized SUPT16H acetylation. Four domains of SUPT16H, including N-terminal domain (NTD), dimerization domain (DD), middle domain (MD), and C-terminal domain (CTD), were cloned in pQCXIP vector and expressed with FLAG tag (**Fig 2A**). These vectors were transfected in HEK293T cells, which were subjected to the reciprocal protein IP assays. Both DD and MD domains from acetylation pull-down were detected by FLAG immunoblotting, but only MD domain from FLAG pull-down was detected by acetylation immunoblotting (**Fig 2B**). These results suggested that only MD domain of SUPT16H is directly acetylated while DD domain likely binds to other acetylated proteins. We further confirmed the functional relevance of TIP60 and BRD4 using the SUPT16H MD domain. TIP60 knockdown significantly reduced the acetylation of MD domain in the reciprocal protein IP assays (**Fig 2C**). BRD4 interacted with MD domain, which was abolished due to TIP60 knockdown (**Fig 2D**). We next mapped the acetylation site of SUPT16H MD domain. Earlier proteomic analyses predicted that K674 of MD domain has the highest likelihood of acetylation (**Fig 2E**). We mutated the lysine to arginine at 674 (K674R) of MD domain, which led to the reduction of MD acetylation (**Fig 2F**) as well as MD interaction with BRD4 (**Fig 2G**). Above all, these results verified that MD domain of SUPT16H is acetylated at K674, which is catalyzed by TIP60 and recognized by BRD4.

**Fig 2.**
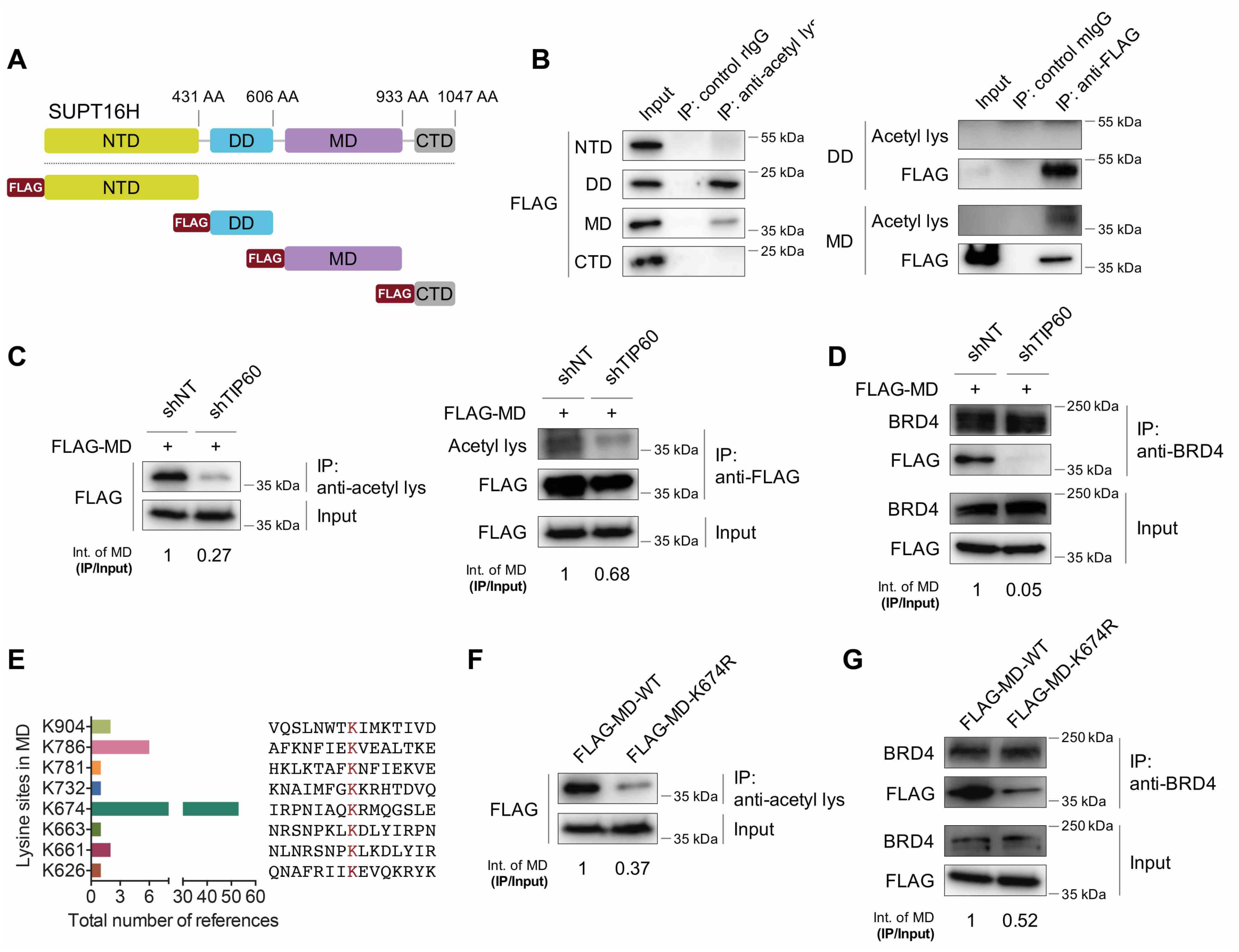
SUPT16H is acetylated at K674 of the middle domain (MD). (**A**) Schematic illustration of defined domains of SUPT16H protein. NTD: N-terminal domain; DD: dimerization domain; MD: middle domain; CTD: C-terminal domain. (**B**) Lysates of HEK293T cells transfected with the vector expressing indicated FLAG-tagged protein domain of SUPT16H were incubated with an acetyl-lysine or FLAG antibody, and immunoprecipitated protein samples were analyzed by protein immunoblotting using a FLAG or acetyl-lysine antibody, respectively. (**C**) HEK293T cells stably expressing TIP60 or NT shRNA were transfected with the vector expressing FLAG-tagged MD of SUPT16H. Cell lysates were incubated with an acetyl-lysine or FLAG antibody, and immunoprecipitated protein samples were analyzed by protein immunoblotting using a FLAG or acetyl-lysine antibody, respectively. (**D**) Lysates of HEK293T cells in (**C**) were incubated a BRD4 antibody. Immunoprecipitated protein samples were analyzed by protein immunoblotting using a FLAG or BRD4 antibody. (**E**) Prediction of acetylated lysine sites in MD of SUPT16H by PhosphoSitePlus®. Lysine site K674 has the highest likelihood to be acetylated. (**F**) Lysates of HEK293T cells transfected with FLAG-tagged, wild-type (WT) or K674R MD of SUPT16H, were incubated with an acetyl-lysine antibody. Immunoprecipitated protein samples were analyzed by protein immunoblotting using a FLAG antibody. (**G**) Lysates of HEK293T cells in (**F**) were incubated with a BRD4 antibody. Immunoprecipitated protein samples were analyzed by protein immunoblotting using a FLAG or BRD4 antibody.

### BRD4 interaction enhances protein stability of SUPT16H

We next addressed what is the functional impact of SUPT16H acetylation. We noticed that treatment of JQ1 causes the notable reduction of SUPT16H protein in all tested cell lines (**Fig 3A**). Consistently, treatment of an alternative BETi UMB-136 (Huang et al., 2017) with the different chemical structure as JQ1 also rendered the similar effect on SUPT16H protein (**Fig 3B**). To reassure it is due to BRD4, we determined the effect of BRD4 knockdown on both mRNA and protein levels of SUPT16H. Our results showed that BRD4 knockdown by its shRNA has no effect on SUPT16H mRNA but leads to the dramatic decrease of SUPT16H protein (**Fig 3C, D**). BRD4 knockdown by its siRNAs yielded the similar results (**Fig 3E, F**). Indeed, the ubiquitination-mediated proteolysis plays an important role in regulating the functions of FACT subunit SUPT16H (Han et al., 2010; Kaja et al., 2021). We further confirmed that treatment of JQ1 strongly increases the K48-linked ubiquitination of SUPT16H, which explains the JQ1-induced reduction of SUPT16H protein likely through the ubiquitination-mediated protein degradation. Taken together, our results delineated that BRD4 binding to SUTP16H prevents K48-linked ubiquitination and protein degradation of SUPT16H.

**Fig 3.**
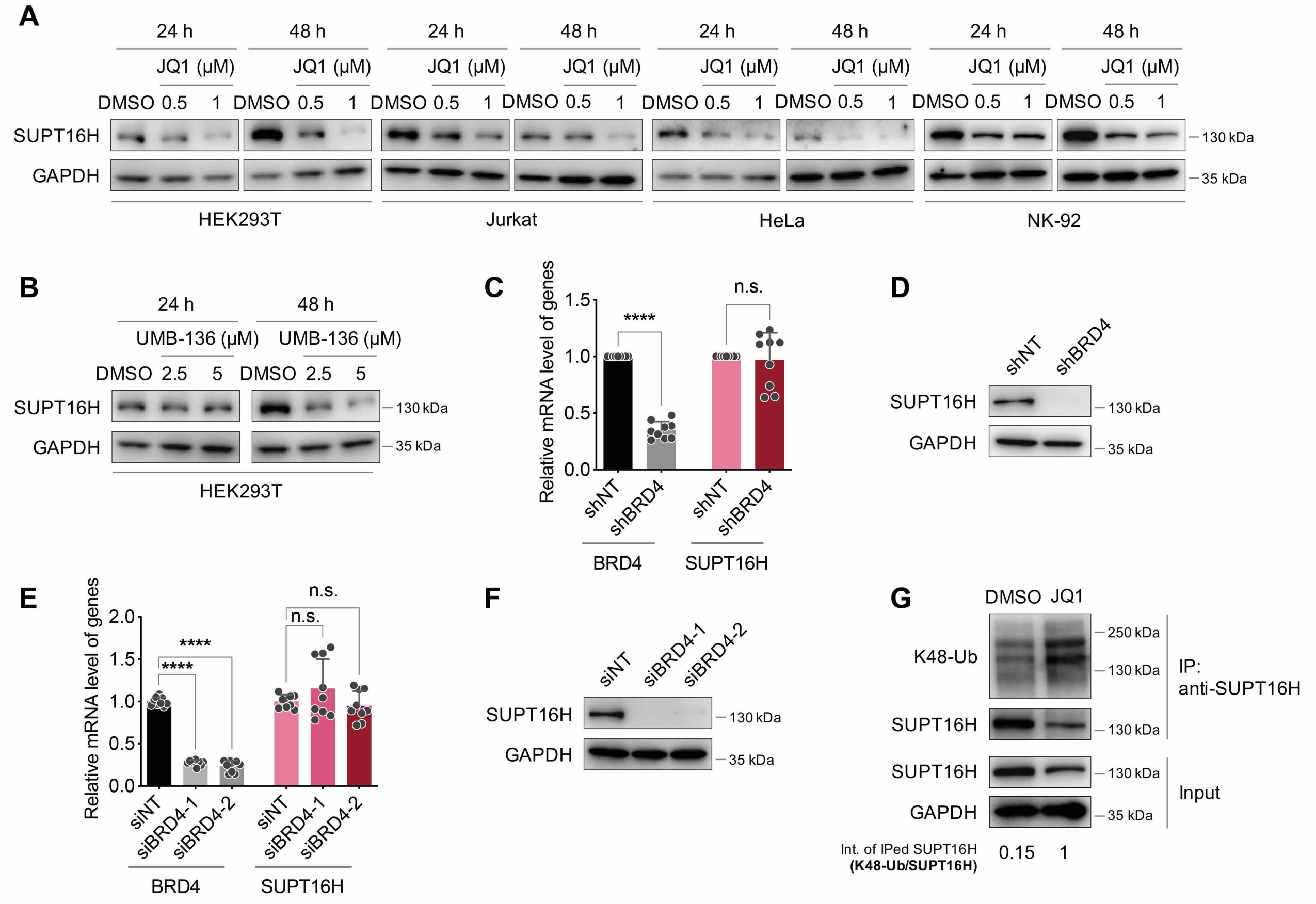
SUPT16H-BRD4 interaction prevents SUPT16H from protein degradation. (**A**) HEK293T, Jurkat, HeLa, and NK-92 cells treated with JQ1 were subjected to protein immunoblotting analysis using a SUPT16H antibody. (**B**) The similar analysis as in (A) was performed for HEK293T cells treated with UMB-136. (**C, D**) HEK293T cells stably expressing BRD4 or NT shRNA were subjected to mRNA RT-qPCR analysis of BRD4 or SUPT16 expression (**C**) or protein immunoblotting analysis using a SUPT16H antibody (**D**). (**E, F**) The similar analysis as in (**C, D**) was performed for HEK293T cells transfected with BRD4 or NT siRNAs. Results were calculated from three independent experiments (**** *P* < 0.0001, Student’s *t* test for **C**, one-way ANOVA for **E**). (**G**) Lysates of HEK293T cells treated with JQ1 were incubated with a SUPT16H antibody. Immunoprecipitated protein samples were analyzed by protein immunoblotting using a K48-ubquitin or SUPT16H antibody.

### SUPT16H-BRD4 binds to epigenetic silencing enzymes leading to gene suppression

Our earlier studies demonstrated that both SUPT16H and BRD4 contribute to the silencing of integrated HIV-1 proviruses (Huang et al., 2015; Zhu et al., 2012), indicating that SUPT16H and BRD4 also possess gene suppression functions, which has not been characterized in details comparing to their well-studied roles in transcriptional activation. We confirmed that knockdown of SUPT16H or BRD4 by their specific siRNAs increases HIV-1 LTR promoter driven gene expression (**Fig S2A, B**). Treatment of the reported SUPT16H inhibitor curaxin 137 (CBL0137) also enhanced HIV-1 LTR promoter activity (**Fig S2C**). Since we identified that BRD4 binds to SUPT16H, we speculated that SUPT16H-BRD4 renders gene suppression functions coordinately.

It has been reported that EZH2 and HDAC1 are two key epigenetic silencing enzymes acting through modulation of histone methylation and deacetylation, respectively. We hypothesized that SUPT16H-BRD4 associates with these epigenetic silencing enzymes contributing to gene suppression. As the supportive evidence, an earlier proteomic study predicted that EZH2 protein is a potential binding partner of SUPT16H (Xu et al., 2015). Our own results confirmed that SUPT16H-BRD4 indeed interacts with EZH2 and HDAC1 by a series of protein IP assays. HEK293T cells were transfected with V5-tagged EZH2 or HDAC1, and the V5 IP led to the pull-down of endogenous SUPT16H and BRD4 (**Fig 4A, B**). Endogenous EZH2 or HDAC1 were IPed using their specific antibodies, which also pulled down endogenous SUPT16H and BRD4 (**Fig 4C, D**). In the reciprocal IP of endogenous SUPT16H, EZH2, HDAC1, and BRD4 were all pulled down as well (**Fig 4E**. **Fig 1F**). Furthermore, we determined the effect of CBL0137 on histone methylation and acetylation marks regulated by EZH2 and HDAC1, H3K9me3, H3K27me3 and H3ac. Treatment of CBL0137 significantly reduced H3K9me3 and H3K27me3 but increased H3ac (**Fig 4F**). These results suggested that one potential new mechanism for SUPT16H-BRD4 to exert gene suppression functions is to interact with epigenetic silencing enzymes (EZH2 and HDAC1) and thus support their activities to reduce chromatin accessibility.

**Fig 4.**
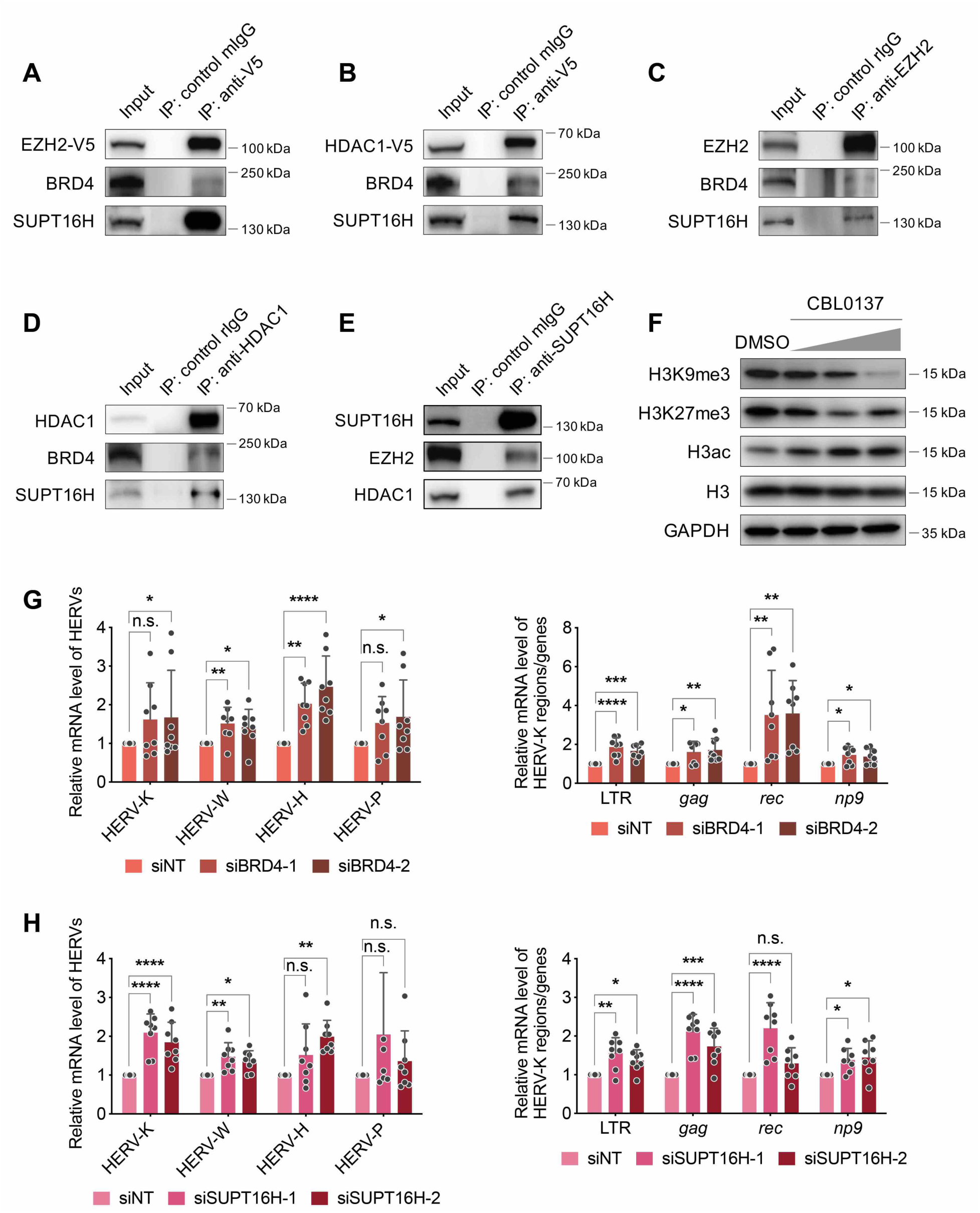
SUPT16H-BRD4 associates with EZH2 and HDAC1 contributing to gene silencing. (**A, B**) Lysates of HEK293T cells transfected with the vector expressing V5-tagged EZH2 (**A**) or HDAC1 (**B**) were incubated with a V5 antibody or mIgG. Immunoprecipitated protein samples were analyzed by protein immunoblotting using a SUPT16H, BRD4, or V5 antibody. (**C-E**) Lysates of HEK293T cells were incubated with a EZH2 (**C**), HDAC1 (**D**), or SUPT16H (**E**) antibody. Immunoprecipitated protein samples were analyzed by protein immunoblotting using the indicated antibodies. (**F**) HEK293T cells treated with CBL0137 at the increasing doses (100, 200, 500 nM) were subjected to protein immunoblotting analysis using a H3K9me3, H3K27me3, H3ac, or histone H3 antibody. (**G, H**) HEK293T cells transfected with BRD4 (**G**) or SUPT16H (**H**) siRNAs were subjected to mRNA RT-qPCR analysis of HERVs expression using primers targeting the *env* gene of HERV-K, W, H, P (left panel), and other regions and genes (LTR, *gag*, *rec* and *np9*) of HERV-K (right panel). Results were calculated from three independent experiments (* *P* < 0.05, ** *P* < 0.01, *** *P* < 0.001, **** *P* < 0.0001, two-way ANOVA).

We further confirmed the gene silencing functions of SUPT16H and BRD4 by using human endogenous retroviruses (HERVs) that closely resemble HIV-1 proviruses as examples. Knockdown of BRD4 or SUPT16H by their specific siRNAs indeed upregulated the expression of several HERVs in HEK293T cells (**Fig 4G, H**). Treatment of CBL0137 also caused the induction of HERVs gene expression in NK-92 cells (**Fig S5A**).

### SUPT16H-BRD4 controls the induction of IFN signaling

Beyond HERVs, we aimed to identify other cellular genes or gene sets subjected to SUPT16H-BRD4 mediated gene silencing. BRD4 has been implicated in regulation of gene expression in IFN signaling, and JQ1 has been reported to induce IFN signaling (Rialdi et al., 2016; Wang et al., 2020). However, such events have not been explored for SUPT16H. Reanalysis of previously published ChIP-seq datasets of SUPT16H and BRD4 (Kolundzic et al., 2018; Najafova et al., 2017) illustrated that their binding profiles near promoter regions of certain cytokines and interferon-stimulated genes (ISGs) are quite similar (**Fig S3A**). Furthermore, we found that knockdown of BRD4 or SUPT16H by their specific siRNAs indeed increases the expression of IFNβ and ISG15 in HEK293T cells (**Fig 5A-C**). Additionally, knockdown or pharmacological inhibition of SUPT16H also caused the upregulation of interleukin (IL) genes, including IL-4, 6, 8, in both HEK293T (**Fig 5D**) and NK-92 cells (**Fig 5E, F**). These results reassured that SUPT16H also participates in the modulation of IFN signaling, which were further investigated. We showed that knockdown of SUPT16H enhances the luciferase expression driven by interferon stimulating responsive element (ISRE) and interferon-gamma activated site (GAS), which mediate the activation of type I and II IFN responses, IFNα/β and IFNγ, respectively (**Fig 5G, H**). Furthermore, treatment of CBL0137 led to the dramatic decrease of local histone inactive marks (H3K9me3, H3K27me3) but increase of active mark (H3ac) near promoter regions of ISGs and ILs in both HEK293T (**Fig 5I, S3B**) and NK-92 cells (**Fig 5J, S3C**). This is consistent with the other findings that treatment of CBL0137 induces the expression of IFNs and ISGs in NK-92 cells (**Fig S5B**). In conclusion, we provided the evidence that FACT subunit SUPT16H suppresses IFN signaling and its inhibitor CBL0137 blocks such effects.

**Fig 5.**
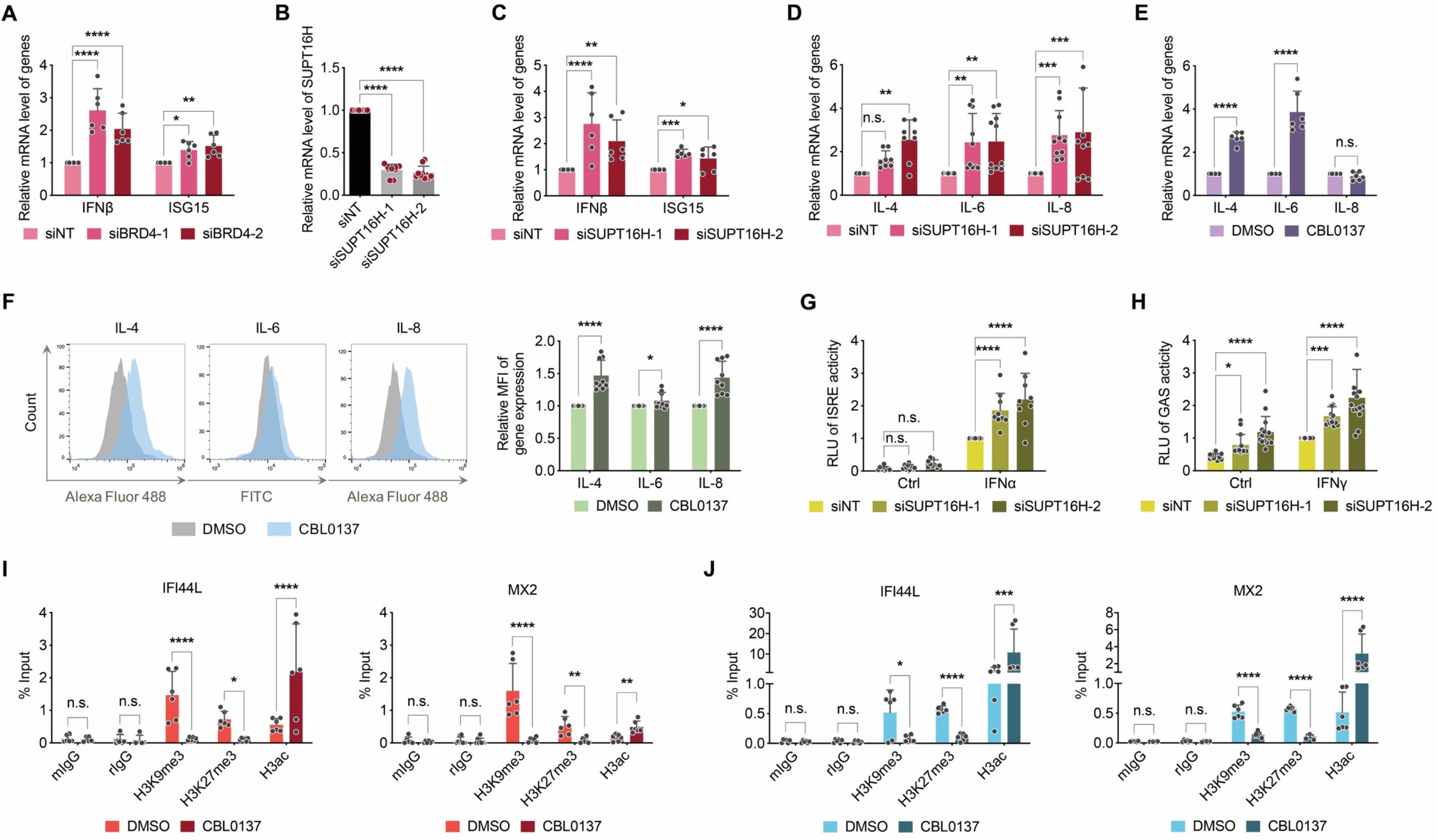
SUPT16H-BRD4 suppresses gene expression of IFN signaling. (**A-C**) HEK293T cells transfected with BRD4 (**A**) or SUPT16H (**C**) siRNAs were subjected to mRNA RT-qPCR analysis of IFNβ and ISG15 expression. Knockdown of SUPT16H by its siRNAs was confirmed by RT-qPCR (**B**). (**D**) HEK293T cells transfected with SUPT16H or NT siRNAs were subjected to mRNA RT-qPCR analysis of IL-4/6/8 expression. (**E, F**) NK-92 cells treated with CBL0137 were subjected to mRNA RT-qPCR (**E**) or protein immunofluorescence (**F**) analysis of IL-4/6/8 expression. (**G, H**) HEK293T cells transfected with SUPT16H or NT siRNAs were further transfected with an IRSE-driven (**G**) or GAS-driven (**H**) firefly luciferase reporter plus TK-driven *Renilla* luciferase control vectors, which were stimulated by Type I (IFNα) or II (IFNγ) IFNs, respectively. RLU (firefly/*Renilla* luciferase activity) from above cells was calculated and normalized to that of siNT-transfected, IFN-treated cells. (**I, J**) Lysates of HEK293T (**I**) or NK-92 (**J**) cells treated with CBL0137 were crosslinked and incubated with a H3K9me3, H3K27me3, H3ac antibody or normal IgG. Immunoprecipitated DNA samples were analyzed by qPCR analysis using primers targeting the promoter region of ISGs (IFI44L, MX2). Results were calculated from three independent experiments (* *P* < 0.05, ** *P* < 0.01, *** *P* < 0.001, **** *P* < 0.0001, one-way ANOVA for **B**, two-way ANOVA for all other results).

### CBL0137 induces IFN signaling and restricts viral infection

IFN signaling plays a central role in host defense against viral infection. Since the SUPT16H inhibitor CBL0137 has the potency to induce IFN signaling, we expected that it can be used as a novel antiviral agent. We first confirmed that knockdown of SUPT16H by its siRNAs (**Fig 6A, B**) or its inhibition by CBL0137 (**Fig 6C**) upregulates the expression of IFN genes in HeLa cells. Particularly, IFNγ, the first characterized and broad-spectrum antiviral cytokine, was significantly induced in CBL0137-treated HeLa cells measured by immunofluorescence and flow cytometry (**Fig 6D**). Induction of selected ISGs (IFI16, MX1, ISG15) in CBL0137-treated HeLa cells was alternatively verified by protein immunoblotting (**Fig 6E**). We next determined the antiviral effect of CBL0137 on various viruses. Interestingly, treatment of CBL0137 efficiently inhibited infection of ZIKV (**Fig 6F**) and influenza A (**Fig 6G**) viruses in HeLa cells. CBL0137 also exerted the inhibitory effect on infection of influenza A in A549 cells (**Fig S4A**). Since SARS-CoV-2 is a newly emerging coronavirus that causes a global threat, we further tested the antiviral effect of CBL0137 on this virus by using the highly reproducible plaque reduction microneutralization (PRMNT) assay (**Fig 6H**). Indeed, treatment of CBL0137 led to the strong inhibition of SARS-CoV-2 infection in Vero E6 cells, comparable to remdesivir (**Fig 6I**). Additionally, JQ1 also exhibited the moderate anti-SARS-CoV-2 effect (**Fig S4B**). Therefore, our results demonstrated that CBL0137, a promising anticancer drug currently in clinical trials, also potently induces IFN signaling and inhibits infections of diverse viruses, including SARS-CoV-2, in epithelial cells.

**Fig 6.**
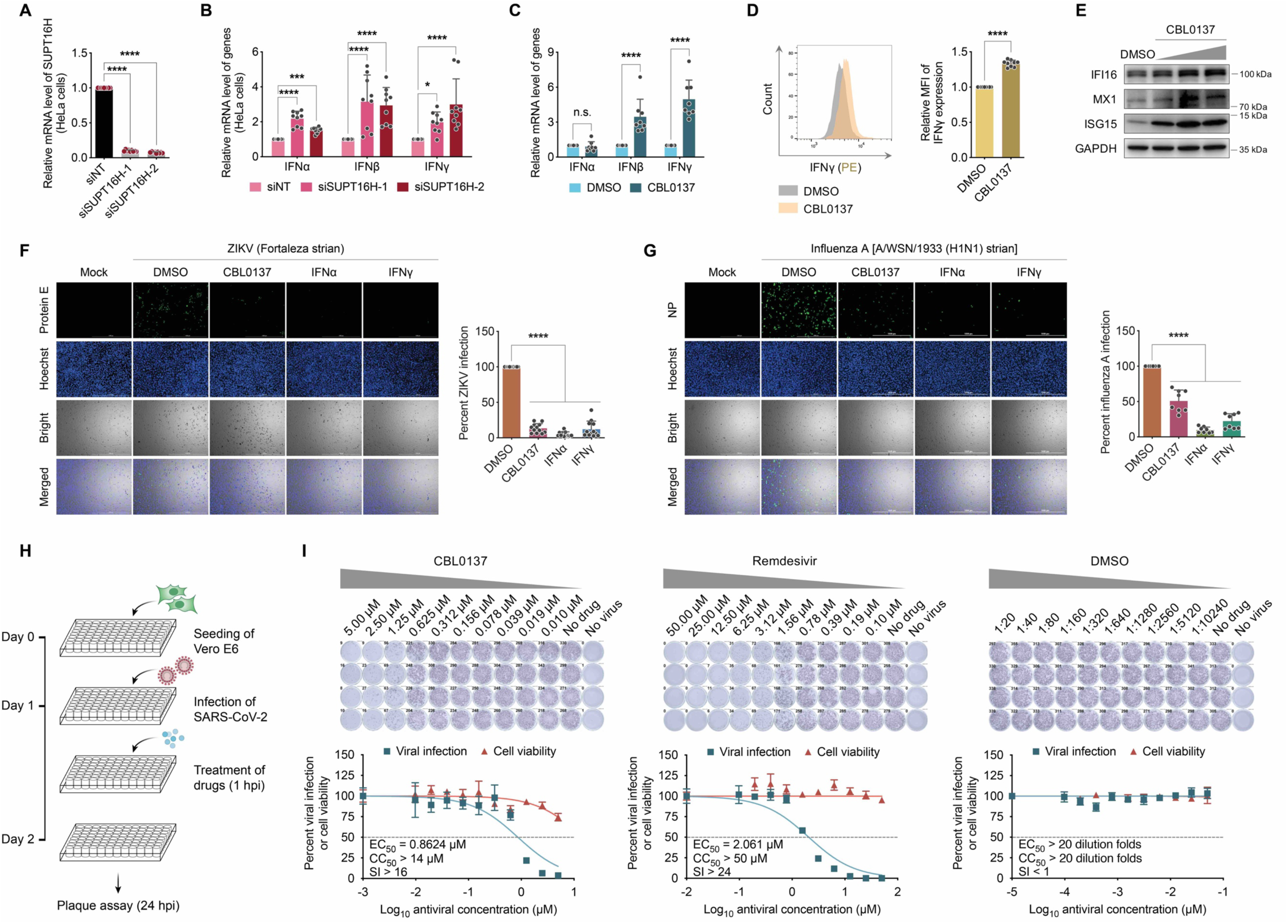
CBL0137 induces IFN signaling and restricts viral infection. (**A, B**) HeLa cells transfected with SUPT16H or NT siRNAs were subjected to mRNA RT-qPCR analysis of SUPT16H (**A**) or IFNs (**B**) expression. (**C**) HeLa cells treated with CBL0137 were subjected to mRNA RT-qPCR analysis of IFNs expression. (**D**) HeLa cells treated with CBL0137 were subjected to protein immunofluorescence analysis of IFNγ by flow cytometry, and results were normalized to DMSO. (**E**) HeLa cells treated with CBL0137 at the increasing doses (100, 200, 500 nM) were subjected to protein immunoblotting analysis of ISGs using the indicated antibodies. **(F, G**) HeLa cells were treated with CBL0137 or IFNs (IFNα or IFNγ), followed by viral infection of ZIKV (**F**) or influenza A (**G**) virus (MOI = 0.5). At 48 hpi (ZIKV) or 24 hpi (influenza A), the above cells were subjected to protein immunofluorescence analysis of viral protein (ZIKV: protein E; influenza A: NP). Viral infection rate was calculated and normalized to DMSO. (**H**) Schematic illustration of PRMNT assay for SARS-CoV-2 infection. (**I**) Vero E6 cells were briefly infected with SARS-CoV-2 (100 – 200 PFU/well), followed by treatment of indicated compounds (CBL0137, remdesivir, or DMSO). At 24 hpi, the above cells were subjected to PRMNT assay at four biological replicates. Results were calculated from three independent experiments (*** *P* < 0.001, **** *P* < 0.0001, Student’s *t*-test for **D**, one-way ANOVA for **A, F, G**, and two-way ANOVA for **B, C**).

### CBL0137 induces NK-mediated killing of virus-infected cells

We demonstrated that SUPT16H acetylation occurs in NK-92 cells (**Fig 1A**). We also observed that treatment of CBL0137 affects gene expression of selected ISGs in NK-92 cells (**Fig S5B**). Thus, we further investigated the impact of the SUPT16H inhibitor CBL0137 on induction of IFN signaling in NK cells. We first performed RNA-seq assays for CBL0137-treated NK-92 cells, which revealed that CBL0137 causes the systemic upregulation of IFNs and ISGs with statistical significance (**Fig 7A**). For selected ISGs upregulated by CBL0137, there was a strong correlation of RNA-seq and RT-qPCR results (**Fig 7B, 7C, S5B**). Pathway analysis confirmed that certain gene sets are enriched from RNA-seq assays of CBL0137-treated NK-92 cells, including responses to virus, IFNγ, and type I IFNs (**Fig 7D**). We also performed the similar analysis for previously published RNA-seq datasets of CBL0137-treated MV4-11 acute myeloid leukemia (AML) cells (Somers et al., 2020), which resulted in the similar finding that CBL0137 significantly induces the expression of cellular genes enriched in IFN signaling (**Fig S6**). This is important, since it indicated that CBL0137 effect on IFN signaling is a universal event independent of cell types. Since IFNs production, especially IFNγ, is a hallmark of NK cell activation, we next determined the impact of CBL0137 on cell killing functions NK cells. In the well-established NK-92 and K562 co-culture assays (**Fig S7**), treatment of NK-92 cells with CBL0137 caused the drastic increase of CD107a and IFNγ expression with the stimulation of K562 cells (**Fig 7E**). Finally, we evaluated the potential of CBL0137 to boost NK cell-mediated killing of virus-infected cells using ZIKV as an example (**Fig 7F**). CBL0137-treated NK-92 cells were co-cultured with ZIKV-infected HeLa cells, which indeed enhanced NK cell-mediated cytotoxicity (**Fig 7G**). More importantly, we observed the similar effect of CBL0137 by using primary NK cells isolated from peripheral blood mononuclear cells (PBMCs) of healthy donors (**Fig 7H**). To summarize, our results demonstrated that beyond the antiviral effect in epithelial cells directly infected with viruses, CBL0137 is also capable of inducing IFN signaling and thus activating NK cells to execute the killing of virus-infected epithelial cells, adding another layer of therapeutic potential of CBL0137 to treat viral infections.

**Fig 7.**
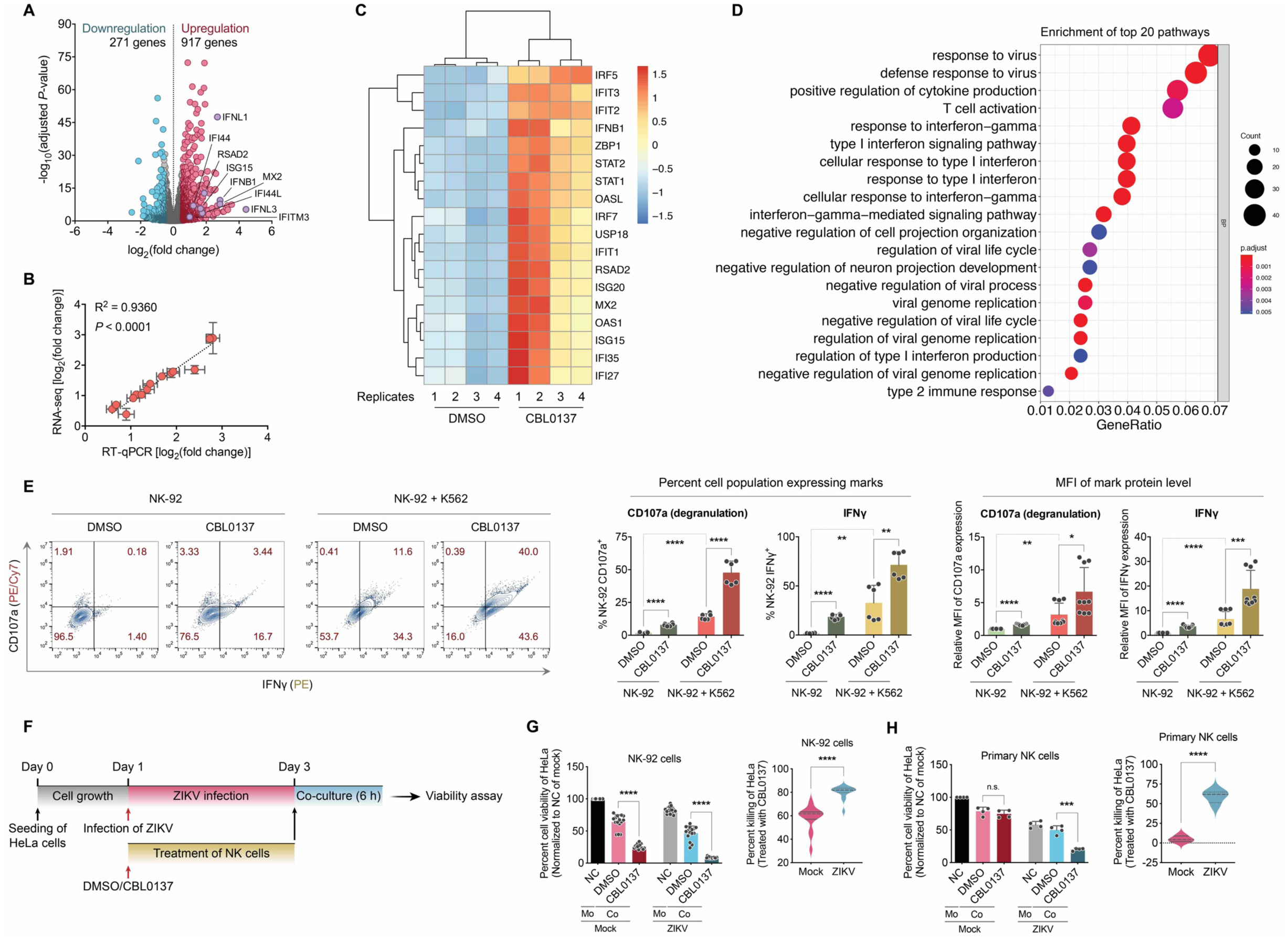
CBL0137 activates NK cell-mediated killing of virus-infected cells. (**A**) Total RNAs were extracted from NK-92 cells treated with CBL0137 or DMSO at four independent replicates, and subjected to RNA-seq analysis. A volcano plot was constructed for RNA-seq results. (**B**) Correlation between RNA-seq and RT-qPCR data for the selected ISGs was determined by Pearson’s *r* with statistical significance. (**C**) Upregulation of selected ISGs in RNA-seq data of NK-92 cells treated with CBL0137 was illustrated in a heatmap (R package: pheatmap). (**D**) Pathway analysis was performed for RNA-seq data of NK-92 cells treated with CBL0137 using GO (R package: clusterProfiler). (**E**) NK-92 cells treated with CBL0137 or DMSO were co-cultured with or without K562 cells, followed by the protein immunofluorescence analysis of CD107a or IFNγ in NK-92 cells by flow cytometry. Percentage of cell population expressing CD107a or IFNγ as well as the mean fluorescence intensity (MFI) of CD107a or IFNγ were calculated and normalized to DMSO without K562 stimulation. (**F**) Schematic illustration of NK cell-mediated killing of ZIKV-infected cells. (**G, H**) NK-92 (**G**) or primary NK (**H**) cells treated with CBL0137 or DMSO, and co-cultured with mock or ZIKV-infected HeLa cells. Viability of HeLa cells was measured by using ATP-based assay (left panel), and converted to cell killing efficacy (right panel, lines in plots indicate median and 25%/75% percentile). Viability of HeLa cells without co-culture with NK cells nor ZIKV infection was set as 100% (NC: negative control; Mo: mono-culture of HeLa; Co: co-culture of HeLa with NK). Results were calculated from four biological replicates for **H**, and three independent experiments for all others (* *P* < 0.05, ** *P* < 0.01, *** *P* < 0.001, **** *P* < 0.0001, two-way ANOVA for **G, H** [left panel], **E**, and Student’s *t*-test for **G, H** [right panel]).

## Discussion

Histone chaperones play the key roles in regulating chromatin dynamics, especially nucleosome turnover and gene expression (Hammond et al., 2017). Different from most histone chaperones, FACT complex targets both H2A-H2B dimer and H3-H4 tetramer to equilibrate the assembly/disassembly of nucleosomes, through extensive interactions with histones and nucleosomal DNAs (Liu et al., 2020). SUPT16H is a key subunit of FACT complex and a large protein that mediates the majority of above protein interactions (Hondele et al., 2013; Liu et al., 2020). There is evidence that FACT complex possesses both positive and negative regulations of gene expression, but FACT-mediated gene silencing function has not been well characterized comparing to its transactivation activity. In this study, we reported a novel mechanism granting FACT the gene suppression function (**Fig 8**). We identified that FACT subunit SUPT16H undergoes acetylation, catalyzed by TIP60, and interacts with the acetylation “reader” BRD4, which prevents the protein degradation of SUPT16H. SUPT16H-BRD4 further associates with epigenetic silencing enzymes, including the “writer” EZH2 and “eraser” HDAC1, which contributes to functional overlaps of SUPT16H and BRD4 to suppress gene expression. Furthermore, our studies recognized that cellular genes involved in IFN signaling are the new gene targets subjected to modulation of SUPT16H-BRD4. At last, we demonstrated that the SUPT16H inhibitor CBL0137 is potent to induce IFN signaling in both epithelial and NK cells, forming two host defense layers against viral infections.

**Fig 8.**
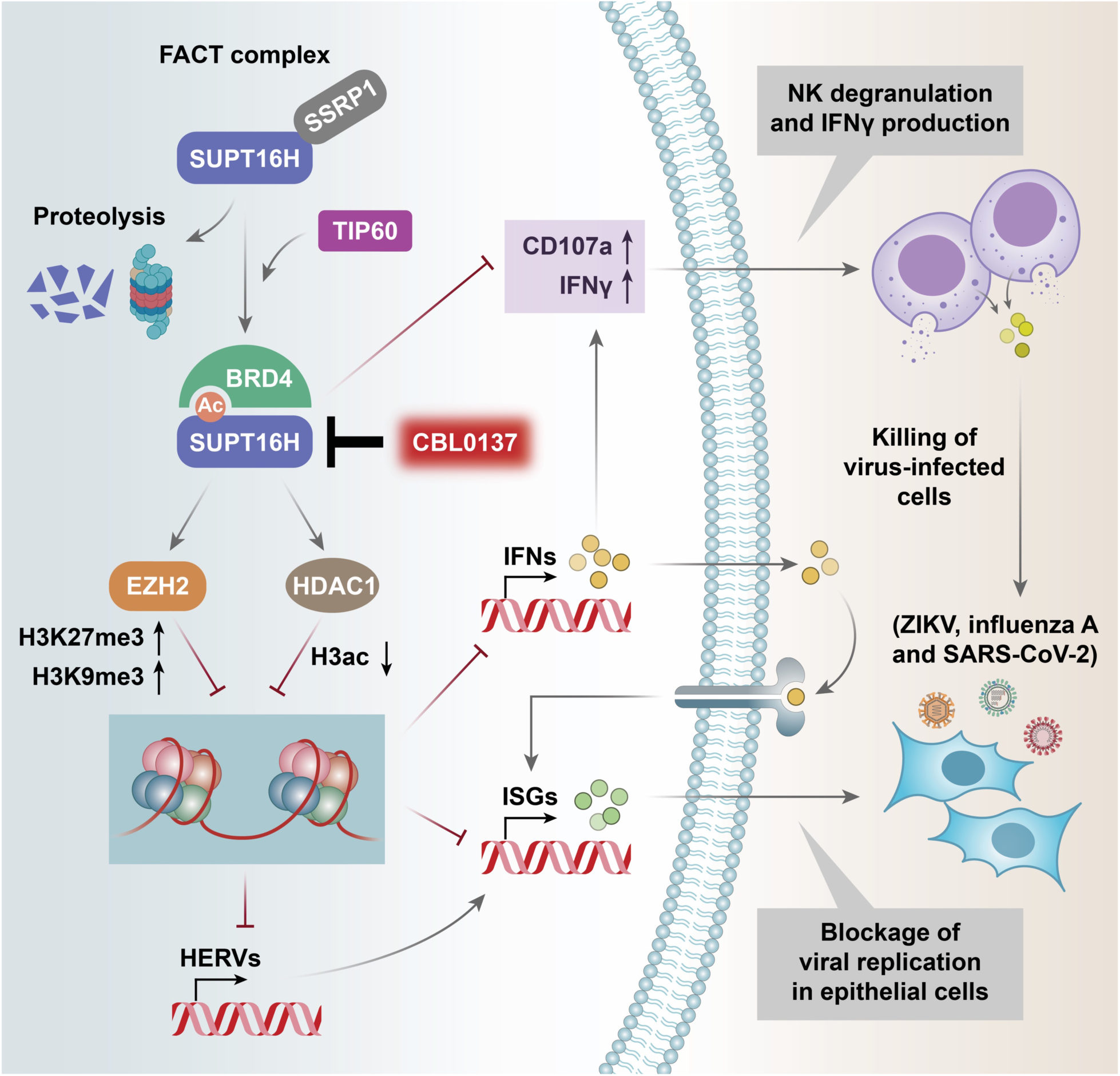
SUPT16H-BRD4 function of gene suppression in antiviral IFN signaling. FACT subunit SUPT16H undergoes protein acetylation at K674 of MD domain, catalyzed by TIP60, which is recognized by BRD4. Such SUPT16H-BRD4 interaction prevents the protein degradation of SUPT16H. SUPT16H-BRD4 further associates with epigenetic silencing enzymes (EZH2, HDAC1) and modulates histone marks (H3K9me3, H3K27me3, H3ac), which overall results in the functional overlaps of SUPT16H and BRD4 to suppress gene expression, including HERVs, IFNs, and ISGs, in multiple types of cells, such as epithelial and NK cells. Furthermore, the SUPT16H inhibitor CBL0137 is potent to induce IFN signaling in both epithelial and NK cells, forming two host defense layers against viral infections.

Although our own data as well as some earlier studies showed that BRD4 is similar as SUPT16H to suppress gene expression in IFN signaling and that treatment of JQ1 induces expression of IFN and ISG genes (Rialdi et al., 2016; Wang et al., 2020), there are also other studies reporting that BRD4 plays an opposite role (Tian et al., 2017). A plausible explanation is that BRD4 likely recognizes other acetylated protein targets, so that its depletion or inhibition may create a more profound impact. BRD4 interaction with acetylated protein targets, including SUPT16H, would rather be dynamic and highly dependent on cellular environment and activation status, which would overall influence BRD4’s function in gene regulation. Thus, BRD4 may disconnect from SUPT16H in terms of their gene suppression function although they interact with each other. Such discrepancy could occur across different cell types and conditions. However, we observed that treatment of CBL0137 causes the fairly consistent impact on inducing IFN signaling in different cell types, indicating that SUPT16H may play a more robust role in silencing IFN and ISG genes than BRD4.

Prior to our studies, there were some reports indicating the potential role of other histone chaperones, but not FACT complex, in regulating IFN signaling (Kadota and Nagata, 2014; McFarlane et al., 2019). Our results provided the new evidence to link FACT complex with silencing of IFN signaling, which extends our knowledge regarding its profound modulatory activities. Our results also bear translational significance by illustrating the new effect of CBL0137 on IFNs induction. Numerous studies have demonstrated the anticancer potential of CBL0137 through regulation of NF-kB pathway (Barone et al., 2017; Gasparian et al., 2011; Kim et al., 2016). Identification of CBL0137’s effect on IFNs activation would further guide its application for treating cancers via immunomodulation. On the contrary, CBL0137’s antiviral potential has never been explored. Our results showed that CBL0137 exerts the antiviral activities in both non-immune (epithelial) and immune (NK) cells, which is worthy of further *in vivo* evaluation, since CBL0137 has already been investigated in animal models of certain cancers (Barone et al., 2017; Carter et al., 2015; Gasparian et al., 2011) and advanced to clinical trials. Particularly, CBL0137 can be considered as an antiviral drug candidate against SARS-CoV-2, since there is so far still no effective antiviral therapy for treating this devastating virus that causes the global pandemic.

## Materials and Methods

### Cells and plasmids

HEK293T, HeLa, Vero E6, and TZM-bl cells were maintained in Dulbecco’s modified Eagle’s medium (Sigma-Aldrich). A549 cells were maintained in Ham’s F-12K (Kaighn’s) medium (Gibco). Jurkat cells were maintained in RPMI 1640 medium (Gibco). K562 human myelogenous leukemia cells were maintained in Iscove’s Modified Dulbecco’s Medium (ATCC). All of above media were supplemented with 10% fetal bovine serum (FBS) (Gibco) and 1 × penicillin-streptomycin solution (Corning). NK-92 (ATCC® CRL-2407™) human natural killer cells were cultured in Alpha Minimum Essential medium according to guideline from ATCC. Human PBMCs and primary NK cells were maintained in RPMI complete medium (15% FBS, 1 × penicillin-streptomycin solution, 1 × MEM Non-Essential Amino Acid Solution, 1 × Sodium Pyruvate, and 20 mM HEPES) supplied with 30 U/ml of human recombinant IL-2 (rIL-2, Roche). Four domains of SUPT16H, NTD, DD, MD and CTD, were cloned in pQCXIP (Clontech) with a N-terminal FLAG tag. Site-specific mutation of K674R was introduced in pQCXIP-FLAG-MD by using the QuikChange Lightning Site-Directed Mutagenesis Kit (Agilent Technologies) following the manufacturer’s instructions. TIP60 shRNA (5’-TCG AAT TGT TTG GGC ACT GAT-3’) and firefly luciferase (FLuc) shRNA (5’-CAC AAA CGC TCT CAT CGA CAA G-3’) were cloned in a pAPM lentiviral vector as previously described (Huang et al., 2019). *cis*-reporting vectors of IFN activity, pISRE-Luc and pGAS-Luc, were purchased (Agilent Technologies). *Renilla* luciferase (RLuc) reporter vector pRL-TK was purchased (Promega). HDAC1 was cloned in pcDNA-DEST40 with a C-terminal V5 tag. pLX317-EZH2-V5 expression vector was acquired from Sigma-Aldrich.

### Human peripheral blood mononuclear cells (PBMCs) and isolation of primary NK cells

Human PBMCs were isolated from healthy peripheral whole blood (STEMCELL Technologies) by using gradient method with the Ficoll-Paque (GE Healthcare), and frozen for later use. Cryopreserved PBMCs were cultured in RPMI complete medium supplied with 30 U/ml of human recombinant IL-2 for three days, and the CD56^+^CD3^−^ NK cells were isolated by using the human NK cell isolation kit (Miltenyi Biotec) following the manufacturer’s instructions.

### Reagents and antibodies

DMSO was purchased from Fisher Scientific. JQ1 were purchased from Sigma-Aldrich. MG149 was purchased from Selleck Chemicals. CBL0137 was provided by Cayman Chemical and purchased from Fisher Scientific. Human IFNα (alpha 2a) and IFNγ were purchased from PBL Assay Science. BD GolgiStop™ protein transport inhibitor was purchased from BD Biosciences. Remdesivir was purchased from AOBIOUS.

The following antibodies were used in this study. Anti-acetyl lysine antibody was purchased from ImmuneChem. Anti-SUPT16H, anti-SSRP1, anti-TIP60, anti-IFI16, anti-MX1, anti-ISG15, anti-GAPDH and normal mouse IgG antibodies were purchased from Santa Cruz Biotechnology. Anti-BRD4 antibody was purchased from Bethyl Laboratories. Anti-FLAG, anti-V5, and normal rabbit IgG antibodies were purchased from Invitrogen. Anti-K48Ub, anti-histone H3, anti-mouse HRP-linked and anti-rabbit HRP-linked antibodies were purchased from Cell Signaling Technology. Anti-HDAC1 antibody was purchased from Novus Biologicals. Anti-H3K9me3, anti-H3K27me3, anti-H3ac (pan-acetyl), and anti-EZH2 antibodies were purchased from Active Motif. PE/Cyanine7 anti-human CD107a (LAMP-1), PE anti-human IFNγ and anti-IL-6 antibodies were purchased from BioLegend. Anti-flavivirus group antigen antibody that probes ZIKV E protein and anti-SARS-CoV-1/2 NP 1C7C7 antibody was purchased from Sigma-Aldrich. Anti-influenza A virus nucleoprotein (NP) antibody was obtained from BEI Resources. Alexa Fluor 488 goat anti-mouse IgG antibody was purchased from Invitrogen. Anti-IL-4, anti-IL-8, and FITC Mouse anti-rat IgG1 antibodies was purchased from BD Biosciences.

### Protein immunoblotting and immunoprecipitation

Protein immunoblotting and immunoprecipitation (IP) were performed as described previously (Zhou et al., 2020). Briefly, total protein was extracted from cell lysates by using 1 × radioimmunoprecipitation assay (RIPA) buffer containing broad-spectrum protease inhibitors. Protein concentrations were measured by BCA assay, followed by electrophoresis and dry electro-transfer. The membrane was blocked with nonfat milk, and incubated with primary and HRP-conjugated secondary antibodies, followed by the incubation with ECL substrate. To determine acetylation, ubiquitination, and other protein binders of the targeted proteins, cell lysates were incubated with the specific antibodies recognizing the targeted proteins or control IgG, followed by the incubation with protein A/G magnetic beads. Beads containing protein immunocomplexes were washed, eluted, and subjected to protein immunoblotting. The intensity of protein bands was quantified by using the ImageJ software.

### Chromatin immunoprecipitation (ChIP)

ChIP was performed as previously described (Huang et al., 2019). In brief, 1% paraformaldehyde (PFA) (Electron Microscopy Sciences) was used for cell cross-linking, followed by the addition of 125 mM glycine to quench the reaction. Cells were then re-suspended by CE buffer and centrifuged to pellet nuclei, which were further incubated with SDS lysis buffer and sonicated to generate DNA fragments. Nuclear lysates were diluted with CHIP dilution buffer and incubated with antibodies recognizing the targeted proteins or control mouse/rabbit IgG, followed by the incubation with protein A/G magnetic beads that were pre-blocked with 0.5 mg/mL BSA and 0.125 mg/mL herring sperm DNA (Invitrogen). The beads were subsequently washed with low-salt buffer, high-salt buffer, LiCl buffer, and TE buffer, and the IPed protein-DNA complexes were eluted by elution buffer. To recover DNA samples, the elutes were treated with 0.2 M NaCl and incubated at 65°C for overnight, followed by the treatment of EDTA, Tris-HCl (pH 6.5), and proteinase K. DNA samples were extracted by phenol/chloroform/isoamyl alcohol (25:24:1). DNA pellets were re-suspended in nuclease-free water, which were used for qPCR analysis. Input (5%) was also included.

### Luciferase reporter assay

HEK293T cells were reversely transfected with gene-specific siRNAs using Lipofectamine™ RNAiMAX Transfection Reagent (Invitrogen). At 72 h post-transfection, cells were further transfected with pISRE-Luc or pGAS-Luc vector along with pRL-TK by using the TurboFect™ Transfection Reagent (Thermo Scientific) for 24 h. Cells were then treated with stimulators (IFNα, IFNγ) for 24 h, followed by the measurement of firefly/Renilla luminescence using the Dual-Glo® luciferase assay system (Promega). Luminescence was measured by the Cytation 5 multimode reader (BioTek), and the relative luciferase unit (RLU) was calculated. To determine the HIV-1 LTR promoter activity, TZM-bl cells harboring LTR-luciferase reporter were either reversely transfected with siRNAs for 72 h or treated with drugs for 24 h, followed by TNFα treatment for 24 h. Luminescence was measured and normalized to total proteins quantified by BCA assay.

### Cell viability assay

ATP-based CellTiter-Glo Luminescent Cell Viability Assay (Promega) was used to measure drug cytotoxicity or cell killing following the manufacturer’s instructions. Luminescence was measured by the Cytation 5 multimode reader (BioTek).

### Immunofluorescence assay (IFA)

IFA for measurement of intracellular proteins followed by flow cytometry analysis was performed as previously described (Zhou et al., 2020). Briefly, 1 × 10^6^ cells were washed with 1 × D-PBS and fixed with 4% PFA. 1 × Perm/Wash buffer (BD Biosciences) containing saponin was used for cell permeabilization, followed by the incubation with primary antibodies and fluorophore-conjugated secondary antibodies. Cell samples were washed and re-suspended with staining buffer, which was subjected to flow cytometry analysis using the BD Accuri C6 Plus with the corresponding optical filters. Mean fluorescence intensity (MFI) was determined by using the FlowJo V10 software.

IFA for measurement of intracellular proteins using microscopy was also performed. HeLa or A549 cells were seeded at the density of 8,000 cells/well on 96-well culture plates. At 24 h post of seeding, cells were treated with CBL0137 (100, 200, or 500 nM), or IFNα (1 × 10^4^ units/ml) and IFNγ (100 ng/ml) for 24 h, followed by the infection of ZIKV (Fortaleza strain) or influenza A virus [A/WSN/1933 (H1N1) strain] (MOI = 0.5). Cells harvested at 48 hpi (ZIKV) or 24 hpi (influenza A virus) were fixed with 4% paraformaldehyde at room temperature (RT) for 10 min and permeabilized with 0.2% Triton X-100 at RT for 15 min, followed by blocking with 1 × D-PBS containing 5% FBS for 1 h at RT. Cells were then incubated with primary antibodies in 1 × D-PBS containing 2.5% FBS for overnight at 4°C, and Alexa Fluor 488-conjugated secondary antibodies for 1 h at RT. Nuclei were stained with Hoechst 33342 (Thermo Scientific) for 10 min at RT. Cells were imaged by using the Cytation 5 multi-mode reader. Percentages of virus-infected cells was quantified by using the Gen5 Image+ software (BioTek).

### Quantitative reverse transcription PCR (RT-qPCR)

Total RNAs were extracted from cells by using the NucleoSpin RNA isolation kit (MACHEREY-NAGEL). Eluted RNA samples were reverse transcribed by using the iScript cDNA Synthesis Kit (Bio-Rad). qPCR was performed by using the iTaq Universal SYBR Green Supermix (Bio-Rad) on a CFX Connect Real-Time PCR System (Bio-Rad). The primers used for qPCR were listed in **Table S1**.

### RNA-seq assay

NK-92 cells were treated with CBL0137 (500 nM) or DMSO for 24 h. RNA samples from four independent repeats were extracted by using the NucleoSpin RNA isolation kit following manufacturer’s manual. RNA samples were submitted to GENEWIZ (South Plainfield, NJ 07080. RNA samples that passed quality control were subjected to rRNA removal through polyA selection, followed by the library preparation. Sequencing was carried out on an Illumina HiSeq platform with the configuration of 2 × 150 bp (paired end), and > 20 million reads per sample were achieved. Differential expression of genes was analyzed by DESeq2 (Love et al., 2014). R packages, pheatmap and clusterProfiler, were used for heatmap construction and pathway analysis, respectively. Published RNA-seq data (GEO accession: GSE126442) (Somers et al., 2020) were analyzed by using the similar methods.

### NK cell cytotoxicity and cell killing assays

NK cell-mediated cytotoxicity was assessed by measuring the expression of degranulation marker CD107a and the production of IFNγ as previously described with slight modifications (Garrido et al., 2018). In brief, K562 target cells were pre-stained with the CellTrace™ CFSE Cell Proliferation Kit (Invitrogen) according to the manufacturer’s protocols. 1 × 10^6^ NK-92 cells were co-cultured with the same number of K562 cells (1 : 1 of effector : target [E : T] ratio) for 4 h. PE/Cy7-CD107a antibody (Biolegend) was added. Cells were then treated with GolgiStop (BD Biosciences) for 1 h, and washed with 1 × D-PBS. After cell fixation and permeabilization, intracellular IFNγ was stained by PE-IFNγ antibody (Biolegend). Fluorescent signal of CD107a or IFNγ in NK-92 cells (CFSE-negative) was determined by flow cytometry analysis. MFI was calculated by using the FlowJo V10 software.

NK cell-mediated killing of virus-infected cells was also performed. HeLa cells were seeded at 24 h prior to viral infection with 8,000 cells/well on 96-well plates. Cells were then infected with ZIKV (MOI = 2) for 48 h. NK-92 or primary NK cells were treated with CBL0137 (500 nM) for 48 h, and CBL0137 was washed away at 24 h post of treatment. NK-92 or primary NK cells were then co-cultured with HeLa cells (2 : 1 or 1 : 5 of effector : target [E : T] ratio) for 6 h. NK cells were removed. HeLa cells were washed twice with DMEM. Survival of HeLa cells was determined by using the ATP-based CellTiter-Glo Luminescent Cell Viability Assay. Percentage of NK cell-mediated killing was calculated as below:

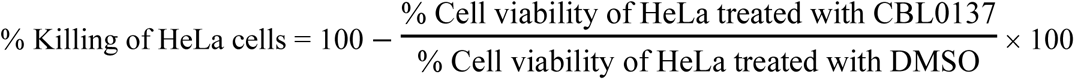

### Plaque reduction microneutralization (PRMNT)

PRMNT assay was performed to evaluate the antiviral activity of drugs against SARS-CoV-2 as previously described (Park et al., 2021). In brief, Vero E6 cells were seeded on 96-well plates with 1 × 10^4^ cells/well at 24 h prior to viral infection. Cells were inoculated with SARS-CoV-2 (USA-WA1/2020 strain) viruses (100 – 200 PFU/well) at 37 °C for 1 h in the CO_2_ incubator. Viral inoculum was removed, and drugs at serial dilutions were added to treat cells for 24 h. Cells were fixed with 10% formalin solution and permeabilized with 0.5% Triton X-100, followed by blocking with 2.5% BSA in PBS. Cells were then incubated with primary antibody targeting SARS-CoV-2 NP and biotinylated secondary antibody. Cells labeled with biotin were detected using the VECTASTAIN® ABC-HRP Kit, Peroxidase (Mouse IgG) (Vector Laboratories) following the manufacturer’s instructions. Viral plaques were counted on the CTL ImmunoSpot plate reader. Infection of wild-type SARS-CoV-2 was carried out at a BSL3 laboratory. Percentage of viral infection was calculated as below:

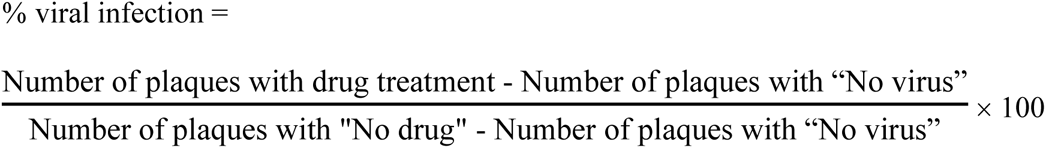

### Statistical analysis

Results were acquired from at least three independent repeats, and analyzed by using either the unpaired, two-tailed Student’s t test or the one-way/two-way analysis of variance (ANOVA). A *P* value less than 0.05 was considered as statistically significant (\**P* < 0.05, \*\**P* < 0.01, \*\*\**P* < 0.001, \*\*\*\**P* < 0.0001). The Pearson’s *r* was used for the correlation analysis. Results were presented as either means ± standard deviation (SD) or means ± standard error of the mean (SEM), and graphed by using the GraphPad Prism 9.0 software.

## Acknowledgments

We thank Wei Zhang (University of Massachusetts at Boston) for providing UMB-136, a BETi derived from 3,5-dimethylisoxazole BETi UMB-32 (McKeown et al., 2014). This study was funded by NIH research grants R01AI150448, R01DE025447, and R33AI116180 to J.Z., and R03DE029716 to N.S.

## Author contributions

J.Z. and D.Z. conceived and designed this study; D.Z. performed most of the experiments; J.G.P. performed the PRMNT assay and its data processing; D.Z., N.S., and J.Z. analyzed the results; Z.W. performed the pathway analysis and heatmap construction of RNA-seq data; J.G.P., Z.W., H.H., G.F., A.B., T.L., Q.M., and L.M.S. contributed materials and/or provided advice for this study; D.Z. and J.Z. wrote the manuscript; J.Z. supervised the entire study.

## Data and materials availability

All data are present in the main paper or as the supplementary materials. All other related data, information, and materials can be acquired through the specific request.

---

**Fig S1.**
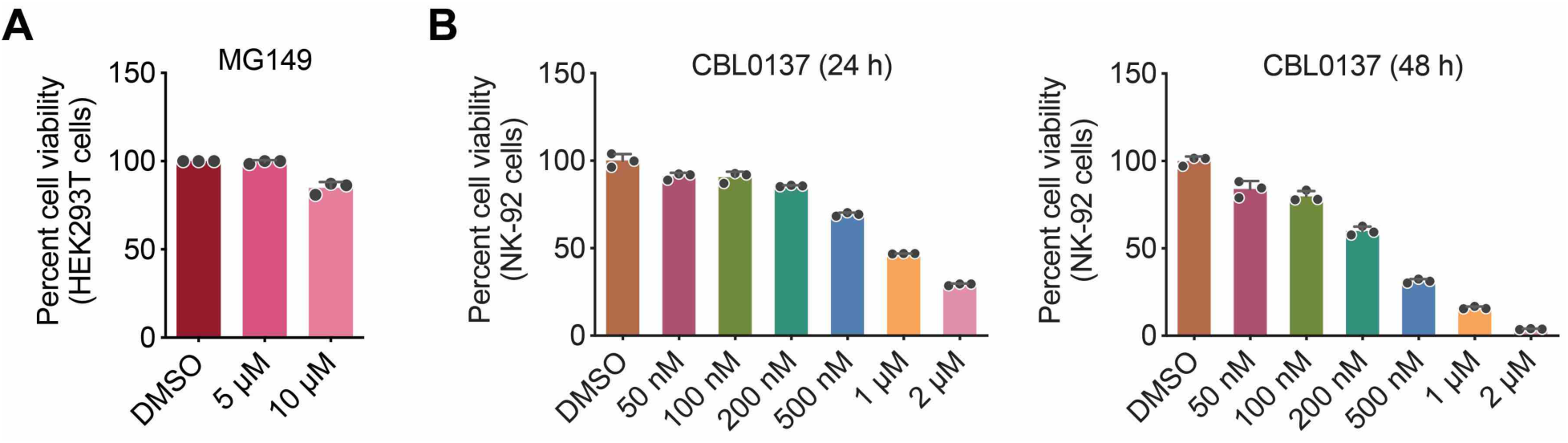
(**A**) Viability of HEK293T cells treated with MG149 was analyzed by ATP-based assay. (**B**) Viability of NK-92 cells treated with CBL0137 was analyzed by ATP-based assay.

**Fig S2.**
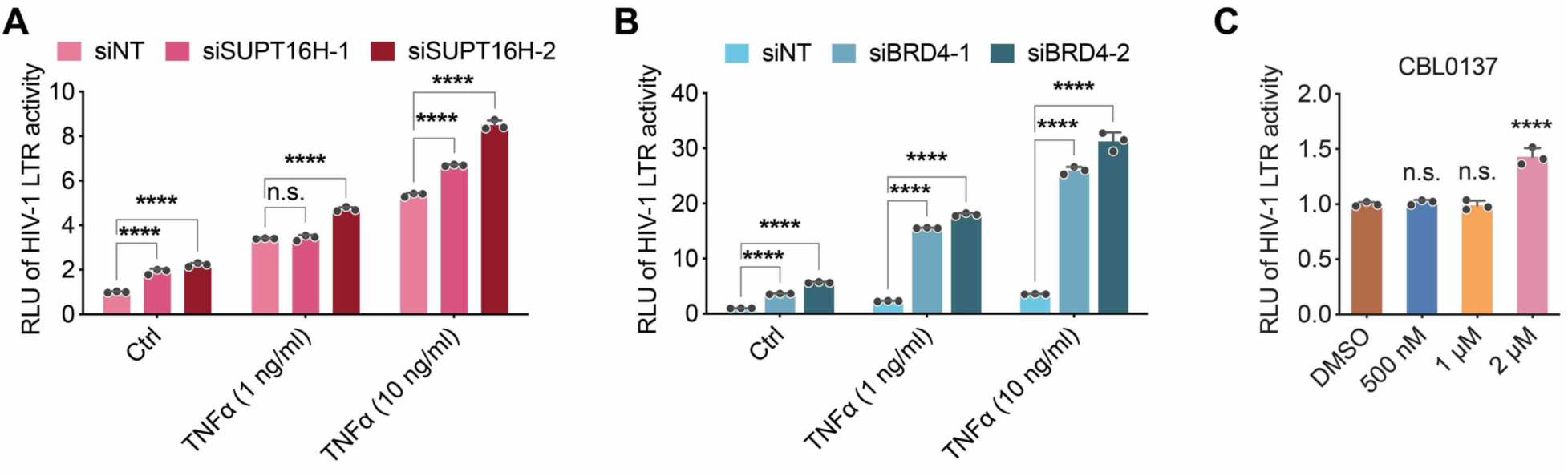
(**A, B**) TZM-bl cells were transfected with SUPT16H (**A**) or BRD4 (**B**) siRNAs, which were then stimulated by TNFα. RLU (firefly luciferase activity/total protein level from BCA assay) from above cells was calculated and normalized to that of siNT-transfected cells without TNFα. (**C**) The similar assay as in (**A, B**) was performed for TZM-bl cells treated with CBL0137. Results were calculated from three independent experiments (**** *P* < 0.0001, two-way ANOVA for **A, B**, one-way ANOVA for **C**).

**Fig S3.**
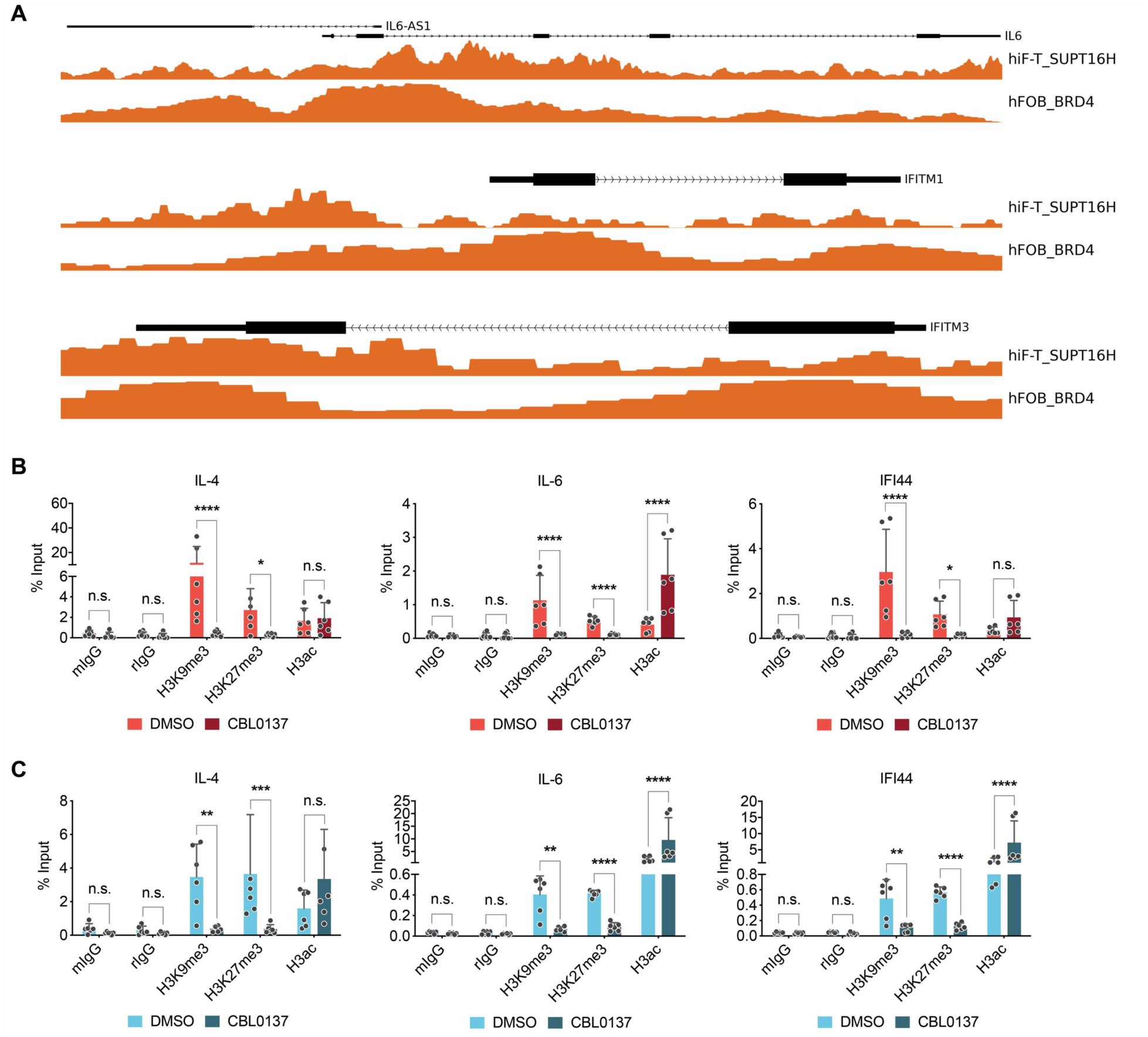
(**A**) Association of SUPT16H and BRD4 at loci of selected ISGs shares the significant overlaps from the analysis of ChIP-seq datasets of SUPT16H and BRD4 (GEO accession: GSE98758 for SUPT16H in hiF-T cells, GSE82295 for BRD4 in hFOB cells). (**B, C**) Lysates of HEK293T (**B**) or NK-92 cells (**C**) treated with CBL0137 were crosslinked and incubated with a H3K9me3, H3K27me3, H3ac antibody or normal IgG. Immunoprecipitated DNA samples were analyzed by qPCR analysis using primers targeting the promoter region of indicated genes. Results were calculated from three independent experiments (* *P* < 0.05, ** *P* < 0.01, *** *P* < 0.001, **** *P* < 0.0001, two-way ANOVA).

**Fig S4.**
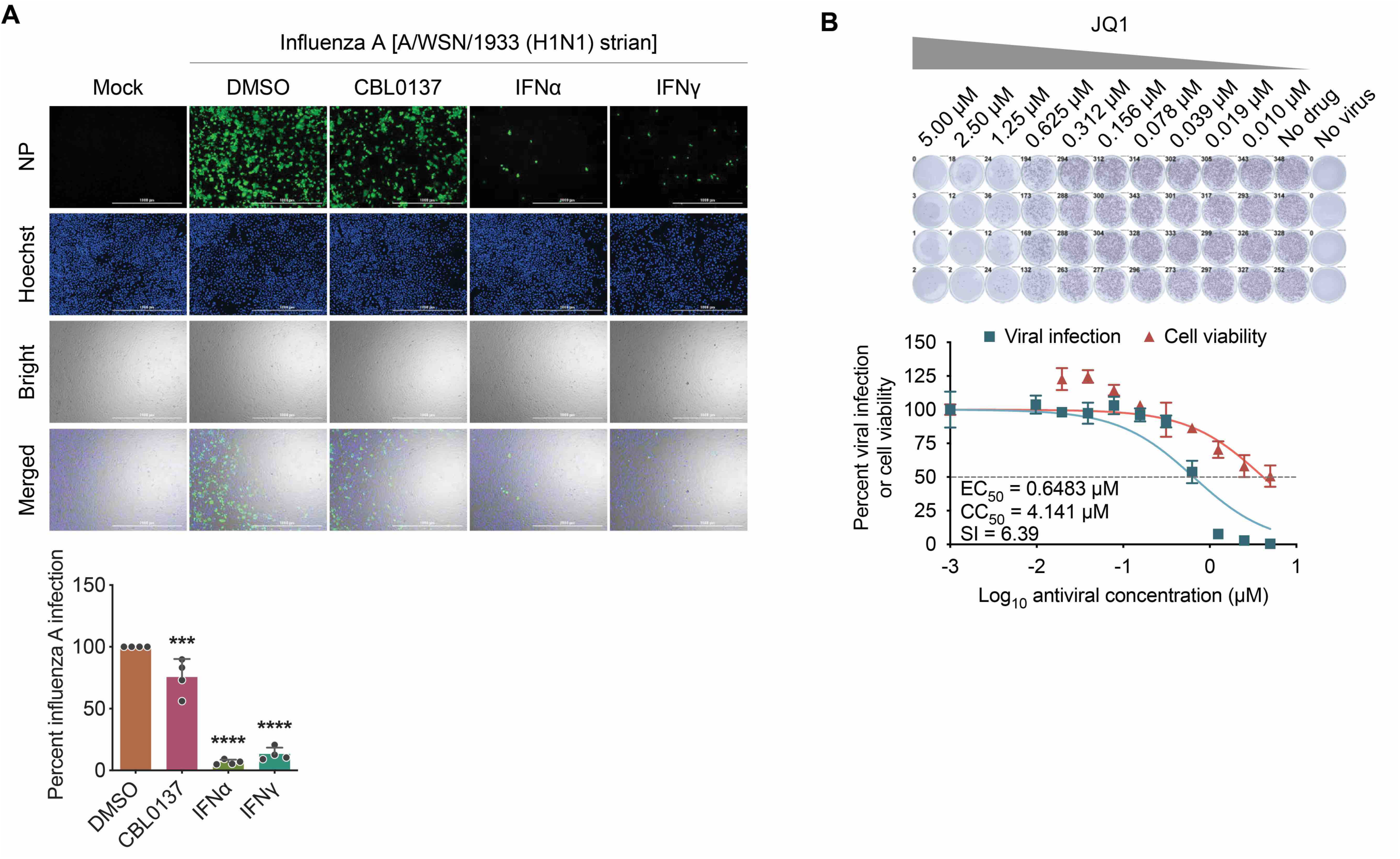
(**A**) A549 cells were treated with CBL0137 or IFN (IFNα or IFNγ), followed by viral infection of influenza A virus (MOI = 0.5). At 24 hpi, the above cells were subjected to protein immunofluorescence analysis of viral protein (NP). Viral infection rate was calculated and normalized to DMSO. (**B**) Vero E6 cells were briefly infected with SARS-CoV-2 (100 – 200 PFU/well), followed by treatment of JQ1. At 24 hpi, the above cells were subjected to PRMNT assay at four biological replicates (*** *P* < 0.001, **** *P* < 0.0001, one-way ANOVA).

**Fig S5.**
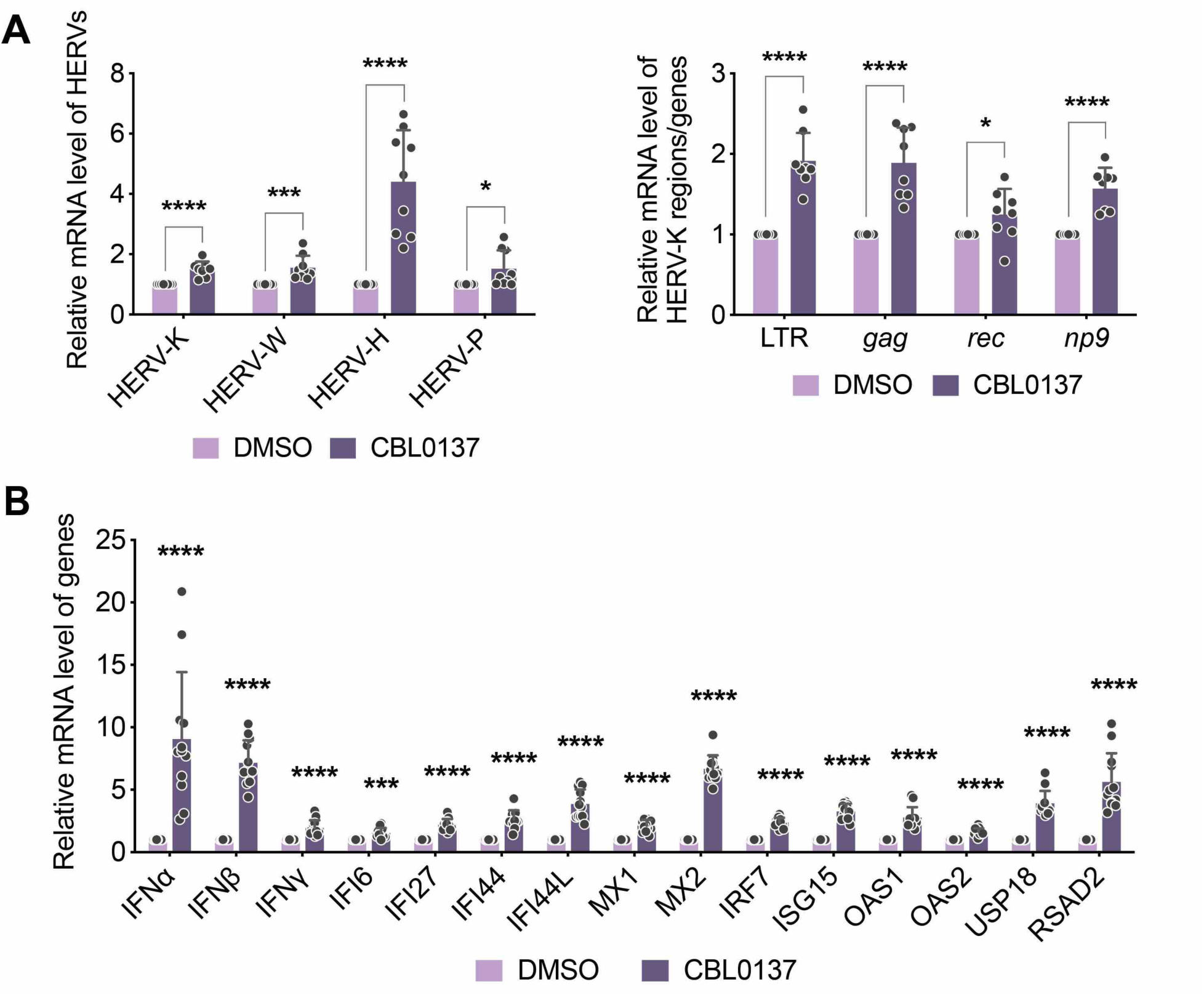
(**A**) NK-92 cells treated with CBL0137 were subjected to mRNA RT-qPCR analysis of HERVs expression using primers targeting the *env* gene of HERV-K, W, H, P (left panel), and other regions and genes (LTR, *gag*, *rec* and *np9*) of HERV-K (right panel). (**B**) NK-92 cells treated with CBL0137 were subjected to mRNA RT-qPCR analysis of indicated IFNs and ISGs. Results were calculated from three (**A**) or six (**B**) independent experiments (* *P* < 0.05, *** *P* < 0.001, **** *P* < 0.0001, two-way ANOVA).

**Fig S6.**
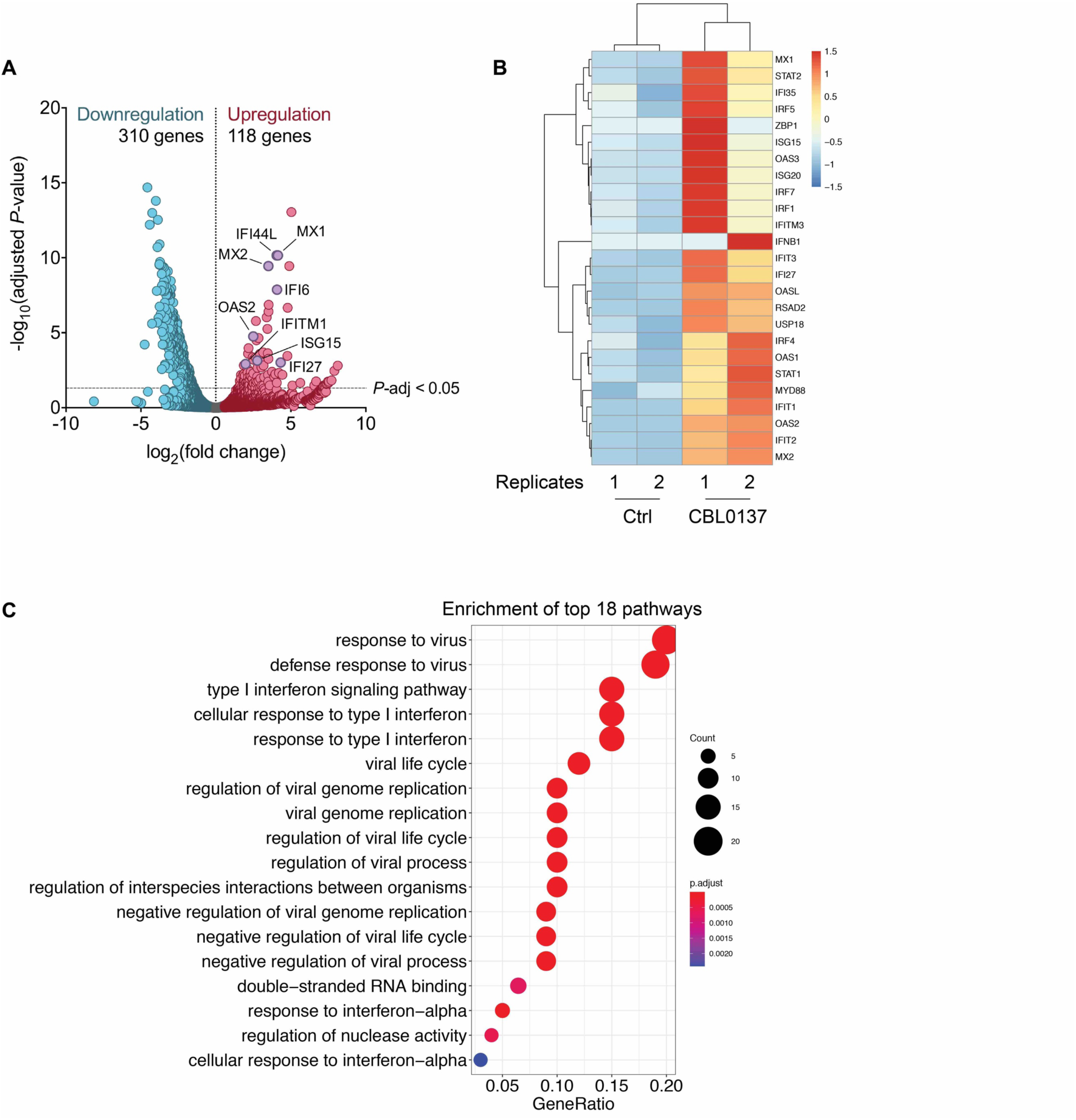
(**A**) A volcano plot was constructed for RNA-seq data (GEO accession: GSE126442) from the earlier study of CBL0137 effect on growth of MV4-11 tumor cells in the NOD/SCID xenograft mice model. (**B**) Upregulation of selected ISGs in the above RNA-seq data was illustrated in a heatmap (R package: pheatmap). (**C**) Pathway analysis was performed for the above RNA-seq data using GO (R package: clusterProfiler).

**Fig S7.**
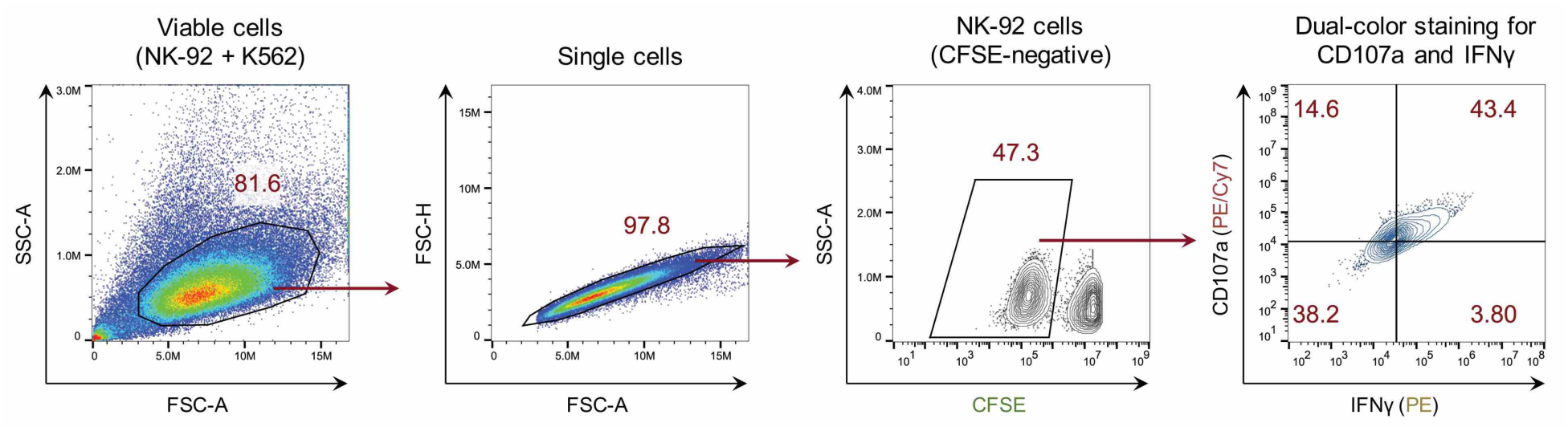
The gating strategy for co-culture assay of NK-92 and K562 cells was illustrated. Viable cells were selected from co-cultured K562 (pre-stained with CFSE) and NK-92 cells by FSC-A vs SSC-A gating. Single cells were selected by FSC-A vs FSC-H gating. NK-92 cells were selected from CFSE-negative population. NK-92 cells were analyzed for expression of CD107a and IFNγ.

**Table S1.**
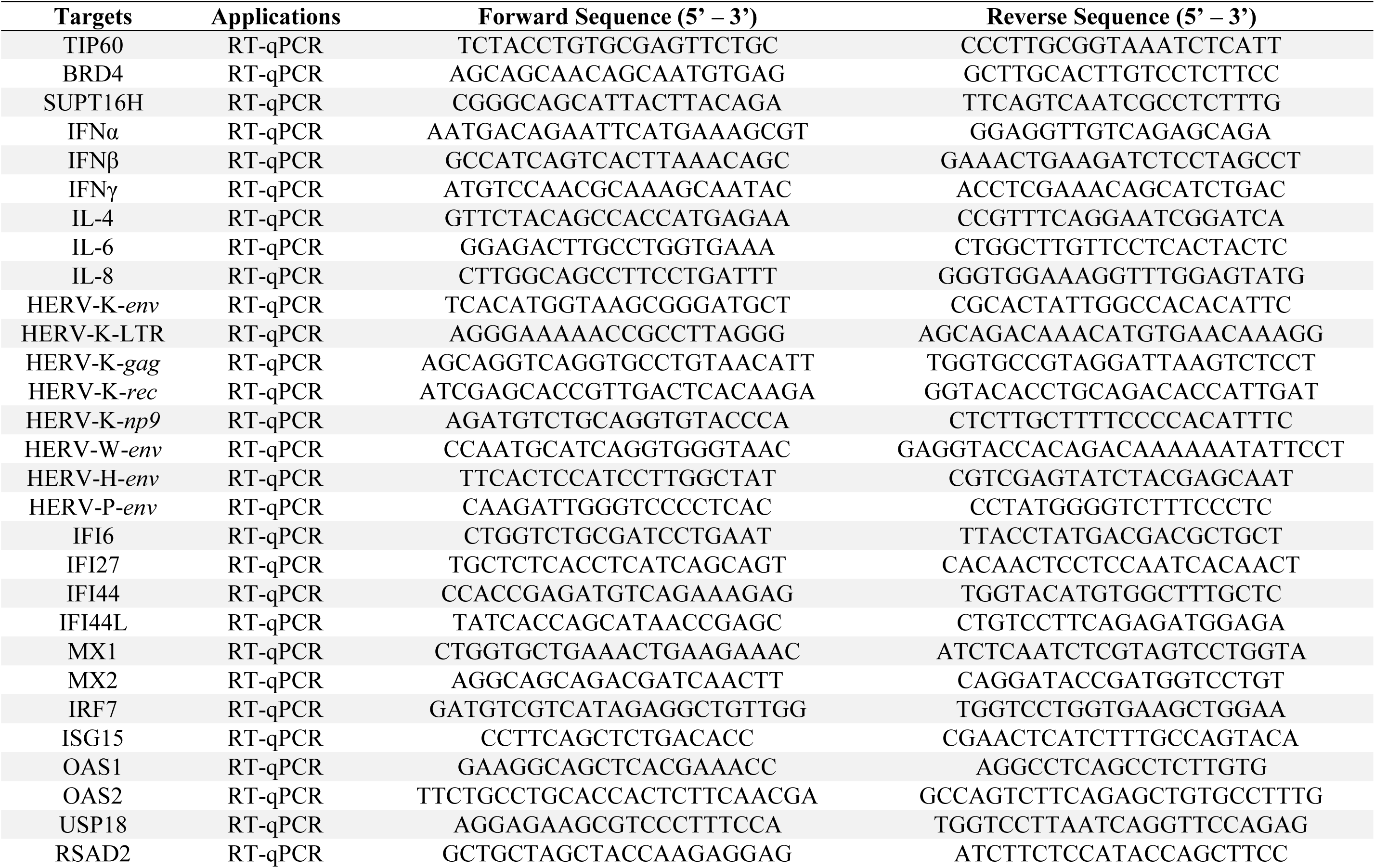
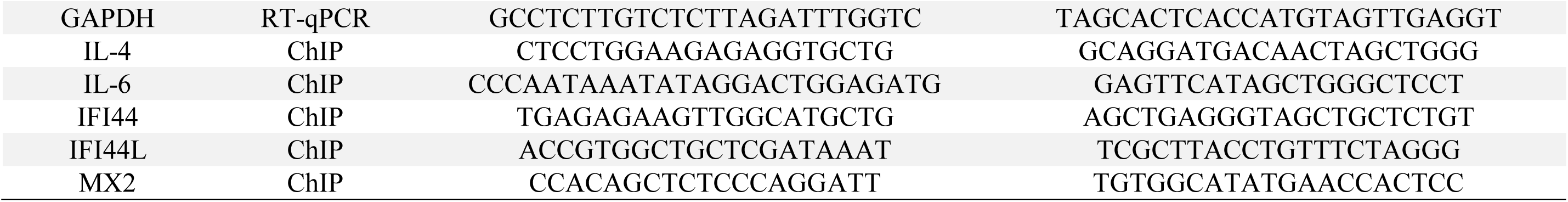
Primers used for qPCR analysis in this study

